# Deciphering the evolutionary history of ectoine catabolism, a compatible solute utilized by *Vibrio diabolicus* as an osmoprotectant and a nutrient source

**DOI:** 10.64898/2026.03.05.709796

**Authors:** Katherine E. Boas Lichty, Heather E. Thomas, Soham M. Bhide, Gary P. Richards, E. Fidelma Boyd

## Abstract

Bacterial adaptation to fluctuations in salinity includes the intracellular accumulation of organic compounds called compatible solutes (CS) such as the amino acid derivatives ectoine and 5-hydroxyectoine. These compounds also play a less appreciated role as readily available nutrients, scavenged from dissolved organic matter in both marine and terrestrial environments. *Vibrio diabolicus* is a marine bacterium originally isolated from deep-sea hydrothermal vents and later shown to have worldwide distribution. In this work, we demonstrated the biosynthesis and uptake of CS ectoine and glycine betaine under high osmotic stress conditions, but not in unstressed *V. diabolicus* cells. A region on chromosome 1 of *V. diabolicus* strain 3098 encoded homologues of genes for ectoine and 5-hydroxyectoine catabolism (*eutDE, ssd_atf_eutBCA)*, regulation (*asnC, enuR*), and transport (ectoine TRAP-type *uehP, uehQM*). Our data showed that ectoine was used as a high energy yielding sole carbon source and the *eutD* gene was essential for ectoine consumption. Phylogenetics based on EutD (DoeA) and gene neighborhood analyses showed that a catabolism cluster was present in *Proteobacteria*, *Thermosulfobacteriota, Bacillota, Actinomycetota,* and *Archaea*. The cluster had a limited phylogenetic distribution in *Gammaproteobacteria* and *Betaproteobacteria* and was widespread in *Alphaproteobacteria*. Phylogenetic reconstruction was consistent with vertical inheritance with gene loss with repeated horizontal acquisitions of the pathway across lineages. The ectoine catabolism pathway was vertically inherited in *Halomonadaceae* and *Vibrionaceae*, with patterns of gene and pathway loss. *Betaproteobacteria Burkholderia, Caballeronia,* and *Paraburkholderia* EutD proteins clustered together and EutD from most *Pseudomonas* species shared a most recent common with this group. EutD from *Alphaproteobacteria* branched in eight divergent clusters with long branch lengths but showed a remarkable conservation of synteny. Catabolism and transporter genes in this group were contiguous and contained either a TRAP-type UehPQM or an ABC-type EhuABCD ectoine transporter. Gram-positive bacteria and *Archaea* were not previously shown to consume ectoines, however, we identified putative ectoine catabolism clusters among *Bacilli, Clostridia, Actinomycetes,* and *Halobacteria*.

**IMPORTANCE:** Ectoine is a well-established CS used to overcome osmotic stress, produced by a wide range of bacteria. The demonstration of ectoine biosynthesis and catabolism in *V. diabolicus* showed that it is conditionally utilized as an osmoprotectant or a nutrient source depending on environmental cues. The conservation of large syntenic blocks of ectoine catabolism, transport, and regulatory genes suggested strong selective pressure to maintain this trait. EutD (DoeA) phylogeny patterns largely followed taxonomy with evidence of horizontal transfer in specific clades, and showed ectoine consumption has a broad taxonomic spread and is lineage enriched. In our dataset, *Alphaproteobacteria* contained the largest diversity of EutD lineages; *Gammaproteobacteria* from marine environments formed a strong secondary group; and EutD from *Betaproteobacteria* were the least diverse. Many species that contained EutD are associated with saline, marine, and plant-associated niches, where ectoine can be scavenged as a nutrient source. The identification of a putative ectoine catabolism pathway in Gram-positive bacteria and *Archaea* needs to be experimentally confirmed and suggests undiscovered diversity to be revealed by future genome sequencing.

## INTRODUCTION

Bacteria have evolved strategies to counteract high osmotic stress, specifically the accumulation of compatible solutes in the cell, which protects vital molecular machinery (1-7). Compatible solutes, also known as osmolytes, are a restricted group of low molecular weight compounds that at high concentrations are compatible with cellular functions and include free amino acids, amino acid derivatives, and quaternary ammonium compounds, amongst others (1-7). Accumulation of osmolytes in the cell is accomplished either by transport from the surrounding environment via specialized transporters or biosynthesis (8-16). The osmolytes ectoine (a cyclic derivative of L-aspartate) and 5-hydroxyectoine (a hydroxylated derivative of ectoine) are used by a wide range of marine bacteria for osmoprotection. The *de novo* biosynthesis of ectoine from endogenous L-aspartate is present in many species (13, 17-21). Ectoine uptake from the surrounding environment for osmoprotection is accomplished using specific transporters, members of the ATP-binding cassette (ABC) family that require ATP, secondary transporters of the **B**etaine **C**arnitine **C**holine **T**ransporter (BCCT) family, and the tripartite ATP-independent periplasmic (TRAP) family of transporters(9-11, 22-28).

Bacteria accumulate compatible solutes in molar concentrations in response to high salinity and once the osmotic stress is removed, these compounds are expelled from the cells (1-7). Thus, osmolytes make up an important component of available dissolved organic matter (DOM) in terrestrial and marine environments that serve as fundamental sources of carbon and nitrogen for growth of heterotrophic bacteria (29-35). The use of ectoines as nutrient sources was first described in *Alphaproteobacteria Sinorhizobium meliloti* and *Ruegeria pomeroyi*, and *Gammaproteobacteria Chromohalobacter salexigens* and *Halomonas elongata* (24, 25, 36, 37). Past studies of the ectoine and 5-hydroxyectoine catabolism pathways showed that ectoine is catabolized to L-aspartate and the enzymes involved were encoded by the *enuR-uehABC-usp-eutABCDE-asnC-ssd-atf* gene cluster in *Ruegeria pomeroyi* (**Fig. 1**) (28, 38). The *eutD* gene encodes a hydrolase that degrades ectoine to N-Acetyl-L-2,4-diaminobutyrate (α-ADABA), which is converted to L-2,4-Diaminobutanoate (DABA) by a deacetylase encoded by *eutE* (**Fig. 2**). The *ssd* and *atf* genes encode enzymes responsible for the conversion of DABA to L-aspartate (**Figs. 1 and 2**) (28, 38). It was later shown that EutD can also convert 5-hydroxyectoine to hydroxy-α-ADABA (39). It was proposed that the *eutABC* genes transform hydroxy-α-ADABA to α-ADABA and hydroxy-DABA to DABA (**Fig. 2**) (39). A tripartite ATP-independent periplasmic (TRAP)-type transporter named *uehABC* is co-transcribed with the catabolism genes in *R. pomeroyi* and was shown to be a high affinity transporter for catabolism (**Fig. 1**) (28). In *R. pomeroyi,* the catabolism genes were inducible in the presence of ectoines, but the metabolic intermediate α-ADABA was the true inducer by binding to the MocR/GabR-type repressor EnuR and relieving catabolic gene repression (39). In *S. meliloti*, the TRAP transporter is substituted for an ABC-type transporter named EhuABCD and was shown to be a high affinity ectoine transporter for catabolism. Similar to *R. pomeroyi*, the catabolism and transporter genes were co-transcribed and induced in the presence of ectoine and 5-hydroxyectoine, but not by NaCl, and also under the control of a EnuR homologue in *S. meliloti* (24, 27). A TRAP-type transporter named TeaABC (*teaABC*) in *H. elongata* participated in ectoine uptake for both osmoprotection and catabolism in this species but did not cluster with the catabolism genes. The *teaABC* genes were induced by NaCl concentration but not by ectoine (11, 36, 40). Kunte and co-workers also demonstrated the presence of ectoine catabolism genes in *Alpha-, Beta-, and Gammaproteobacteria* (36). Bremer and colleagues followed these studies up by examining the evolutionary history of ectoine metabolism, finding that catabolism genes were confined to *Proteobacteria* and proposing an origin within *Alphaproteobacteria* (28, 39, 41). A study examining metabolite metabolism pathways among marine bacteria from the MarRef database (42) suggested that ectoine catabolism is more widespread, based on the presence of EutD and EutE (43).

**Fig. 1.**
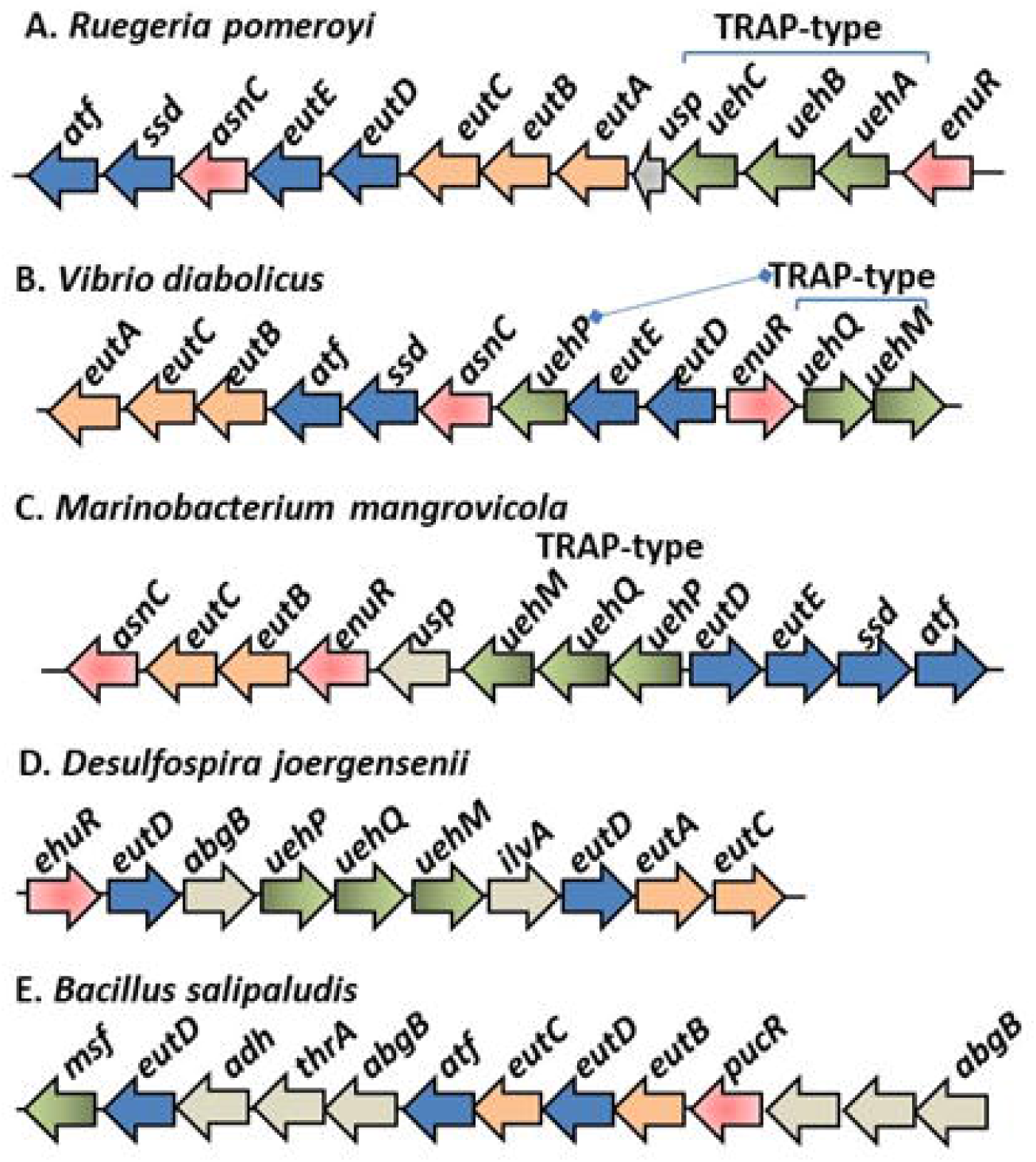
Ectoine and 5-hydroxyectoine catabolism gene clusters. A. Ruegeria pomeroyi, B. Vibrio diabolicus, C. Marinobacterium mangrovicola, D. Desulfospira joergensenii, E. Bacillus salipaludis. Orange arrows represent eutBCA, and blue arrows represent eutDE_ssd_atf. Green arrows represent transporter genes, red arrows indicate regulators (asnC, enuR), and grey ORFs represents proteins whose function in catabolism is unknown.

**Fig. 2.**
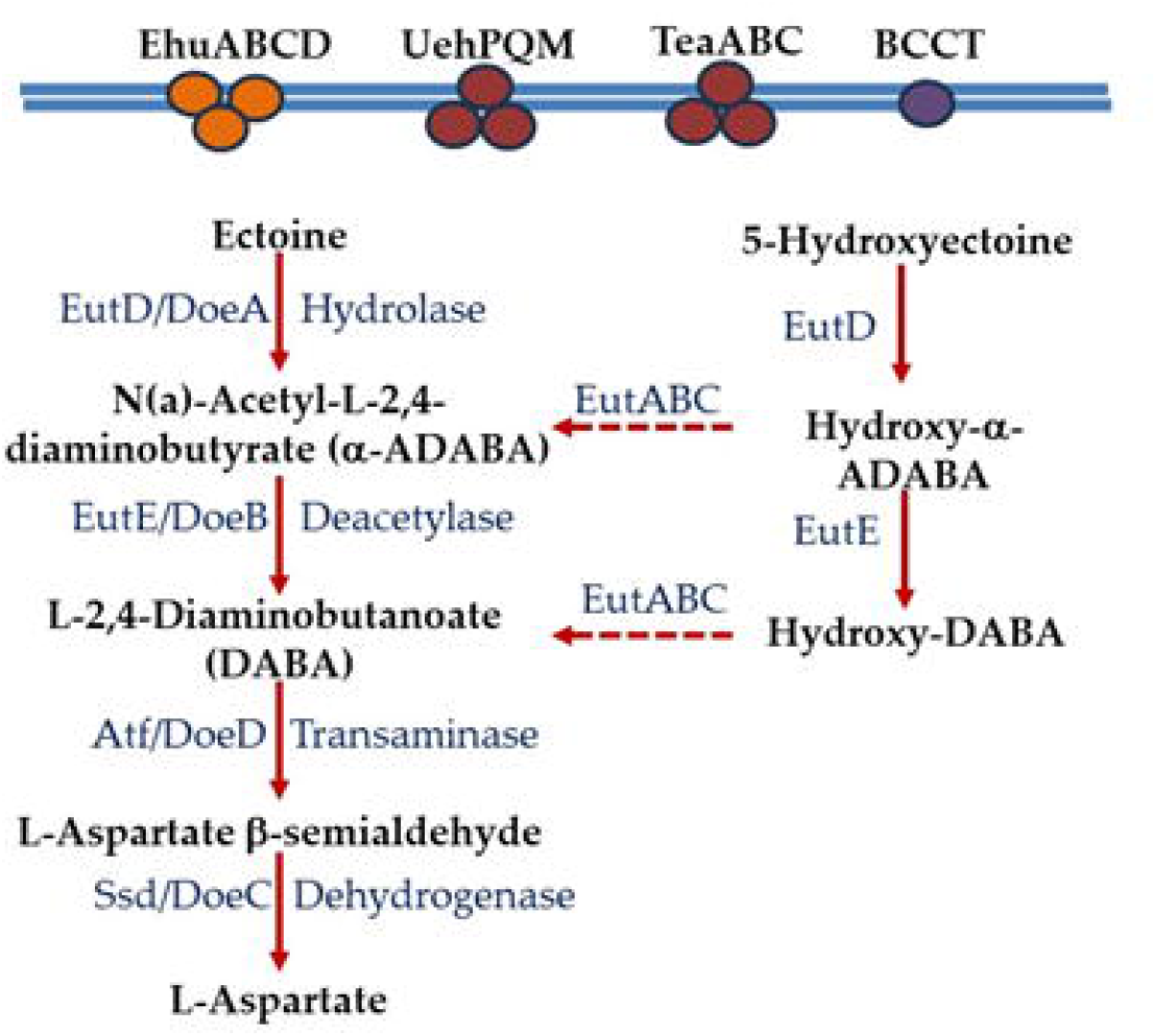
Pathway of ectoine and 5-hydroxyectoine catabolism. The catabolism pathway presented is based upon studies in *Ruegeria pomeroyi*. The enzymes and intermediate compounds in the catabolism pathway as described by Bremer and colleagues (28, 38, 39). Dotted arrows are proposed roles. Double blue line represents inner membrane with ABC-type, TRAP-type and BCCT transporters experimentally shown to uptake ectoine (9-11, 22-28).

*Vibrio diabolicus* is a *Gammaproteobacteria* belonging to the Harveyi clade, first isolated from deep sea hydrothermal vents (44, 45). In the present study, the salinity tolerance of another strain of *V. diabolicus* (strain 3098) (18) was examined by growth pattern analysis at two temperatures across a range of NaCl concentrations. Examination of the genome of *V. diabolicus* 3098 identified glycine betaine (GB) and ectoine compatible solute biosynthesis systems and putative compatible solute transporters. Biosynthesis and uptake of compatible solutes were determined. The *V. diabolicus* genome contained a region on chromosome 1 comprising 12 genes encoding homologues of genes involved in ectoine catabolism, regulation, and transport. The role of this region in ectoine catabolism was examined using growth studies and genetic analysis. The evolutionary origins of ectoine catabolism in *Vibrio* and marine bacteria were determined based on analysis of the EutD protein phylogeny and gene neighborhood analysis.

## RESULTS AND DISCUSSION

### *Vibrio diabolicus* 3098 responses to salinity and temperature

The ability of *V. diabolicus* 3098 to grow in a range of NaCl concentrations was examined by growth patterns in M9 minimal media supplemented with glucose (M9G) at 30°C and 37°C at 0 to 7% NaCl (sea water has an average salinity of 3.5%) (**Fig. S1**). *Vibrio diabolicus* grew at a maximum growth rate of 0.90 h^-1^ at 37°C in M9G 3% NaCl with a final optical density (OD_595_) of 0.72 (**Fig. S1B and Table S2**). Growth of *V. diabolicus* was completely abolished at 37°C in 7% NaCl and at 37°C in the absence of NaCl. At 30°C, *V. diabolicus* grew in the absence of NaCl, but showed poor growth with an OD and growth rate of 0.14 and 0.14 h^-1^ (**Fig. S1A and Table S2**). At 30°C, growth was optimal in M9G 3% NaCl with a final OD of 0.63 and maximum growth rate of 0.60 h^-1^ (**Fig. S1A and Table S2**).

Bioinformatics analysis using homologues of systems previously identified in *Vibrio* species, uncovered the operon *ectABC-aspK* for the *de novo* biosynthesis of ectoine and the operon *betIBA* for GB biosynthesis from its choline precursor (10, 13, 17, 46, 47). An *ectD* homologue required for the biosynthesis of hydroxyectoine from ectoine was not present in *V. diabolicus*. Nine putative compatible solute transporters were identified, two ABC-type transporters, which we named ProU1 (*proVWX*) and ProU2 (*proVWX*), and seven BCCT-type transporters based on similarity to those previously characterized in *Vibrio* species (10, 13, 17, 46, 47). To determine whether *V. diabolicus* can biosynthesize both ectoine and glycine betaine, ^1^H-nuclear magnetic resonance (NMR) spectra analysis was utilized. To accomplish this, ethanol extracts of *V. diabolicus* grown overnight in M9G 5% NaCl were analyzed. Ethanol extracts of unstressed cells, which were cells grown in M9G 1% NaCl, were used as a control and ^1^H-NMR spectra analysis demonstrated the absence of ectoine, L-glutamate, and GB (data not shown). *Vibrio diabolicus* cells grown overnight in M9G 5% NaCl were analyzed and demonstrated the presence of ectoine and L-glutamate (**Fig. 3A**). Furthermore, ethanol extracts of *V. diabolicus* grown in M9G 5% NaCl with the addition of 1 mM choline showed spectra peaks for glycine betaine and the absence of ectoine (**Fig. 3B**). These results are in line with the analyses of other *Vibrio* species in the Harveyi clade that demonstrated the ability to biosynthesis ectoine and GB under osmotic stress conditions (8, 13, 17, 47, 48). Neither *S. meliloti* nor *R. pomeroyi* biosynthesize ectoine whereas *H. elongata* can using the *ectABC_asp* operon.

**Fig. 3.**
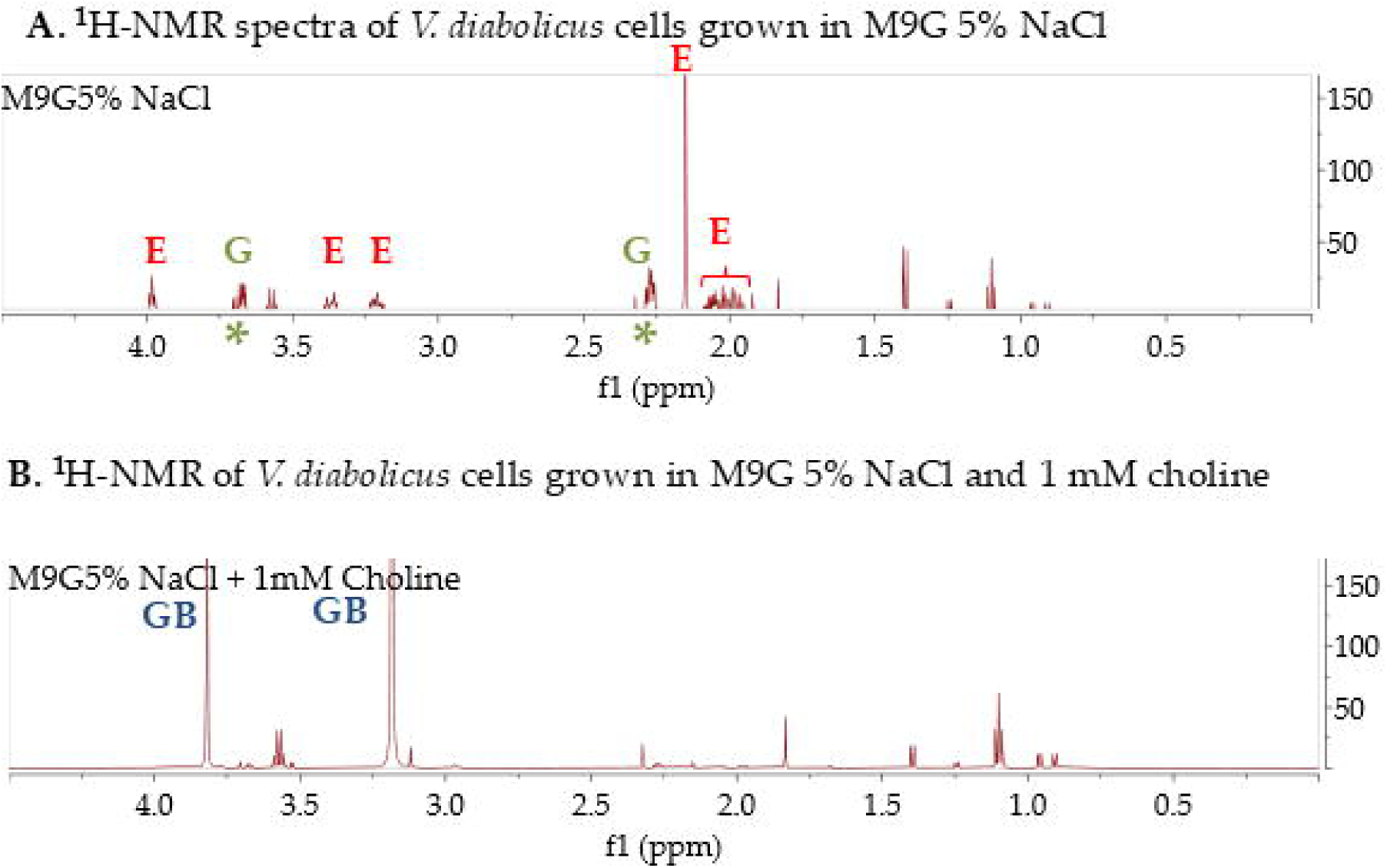
*Vibrio diabolicus* 3098 biosynthesis of compatible solutes ectoine and glycine betaine in response to high NaCl stress. ^1^H-NMR spectra of *V. diabolicus* cellular extract grown in minimal media with 20 mM glucose (M9G) in A. 5% NaCl, and B. 5% NaCl with the addition of 1 mM choline. The spectral peaks corresponding to ectoine (E), glycine betaine (GB), and glutamate (G) are labeled.

The ability of *V. diabolicus* to import compatible solutes in response to high NaCl stress was examined next. To accomplish this, cells were inoculated in M9G 7% NaCl supplemented with GB, dimethylsulfoniopropionate (DMSP), or ectoine and grown for 24 h at 37°C (**Fig. 4**). *Vibrio diabolicus* does not grow in M9G 7% NaCl in the absence of compatible solutes (**Fig. 4**), but cell growth was rescued in the presence of each compatible solute evaluated. M9 media with 1 mM DMSP showed the most efficient growth, followed by GB, and then ectoine (**Fig. 4**). It is of interest to note that both GB and DMSP had a lag phase of ∼3h, whereas ectoine had a lag phase of 9 h. Previous studies in *V. parahaemolyticus, V. natriegens,* and *V. coralliilyticus* also showed uptake of these compatible solutes in high NaCl conditions and GB, DMSP and ectoine all showed similar short lag phases (10, 46, 47). The long lag phase for ectoine in *V. diabolicus* may reflect differences in transporter efficiencies and/or that ectoine is not an essential compatible solute for this species.

**Fig. 4.**
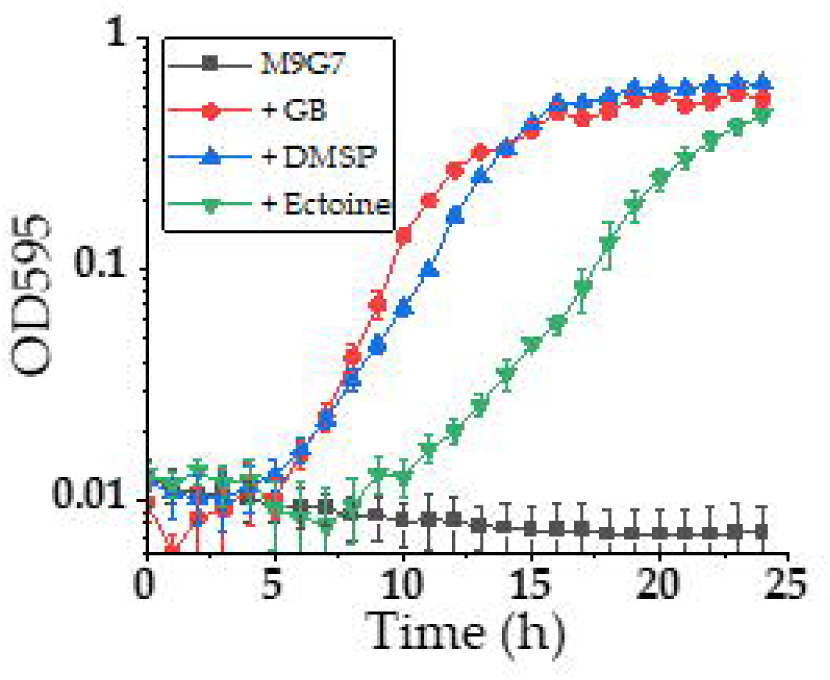
*V. diabolicus* 3098 compatible solute uptake in response to high NaCl stress. Growth curve analysis of *V. diabolicus* 3098 in M9 Glucose and 7% NaCl at 37°C with 1 mM GB (glycine betaine), DMSP (dimethylsulfoniopropionate) or ectoine.

### Ectoine and 5-hydroxyectoine catabolism gene cluster presence in *V. diabolicus*

Genome analysis of *V. diabolicus* 3098 identified homologues of genes involved in catabolism of ectoine and 5-hydroxyectoine to form L-aspartate (**Fig. 1B and Table S1**). A 12-gene cluster with two divergently transcribed regions: *eutDE-uehP-asnC-ssd-atf-eutBCA* and *enuR-uehQM* encoding putative proteins for ectoine and 5-hydroxyectoine catabolism, regulation, and transport was identified (**Fig. 1**). These genes were present in > 90 *V. diabolicus* strains in the NCBI genome database, indicating this gene cluster is ancestral to the species. Sequence homology to the ectoine and 5-hydroxyectoine catabolism, transporter, and regulatory genes from *R. pomeroyi* DSS-3 and *H. elongata* DSM2581 was used to determine gene designation in *V. diabolicus* (28, 36, 38, 49) (**Fig. 1 and Table S1**).

The genes *eutDE* encoded homologues of a hydrolase and deacetylase needed for the first two steps of ectoine to DABA degradation (**Fig. 2**). Next to these genes in *V. diabolicus* was a gene encoding a homologue of the TRAP-type transporter binding protein *dctP*, which we named *uehP* (**Fig. 1 and Table S1).** The *asnC* gene encoded a protein that is a member of the feast or famine regulatory proteins and was found to be essential for the ability of *R. pomeroyi* to use ectoine as a carbon source (28, 36, 38, 50) (**Table S1**). Downstream of this gene were homologues of *ssd* and *atf* that encode enzymes responsible for the conversion of DABA to L-aspartate (**Fig. 1**, **Fig. 2**) (28, 38). Also present was the cluster *eutABC* proposed to be involved in the degradation of hydroxy derivatives in *R. pomeroyi* (**Fig. 2)** (39). *Vibrio diabolicus* contains a GabR/MocR family regulatory protein named *enuR* (ectoine nutrient regulator), shown to be a repressor of ectoine catabolism and transport in *R. pomeroyi* and *S. meliloti* (20, 51). Homologues of TRAP transporter genes *dctQM* were present which we named *uehQM* genes to avoid confusion with ABC-type transporter genes. These genes were separated from *uehP*, and encoded proteins that shared 42%, 44%, and 67% amino acid identity to TRAP-type proteins UehABC respectively from *Alphaproteobacteria R. pomeroyi*, and 41%, 38%, and 64% identity to TRAP-type TeaABC respectively from *Gammaproteobacteria H. elongata* (**Table S1**).

### *Vibrio diabolicus* 3098 utilizes ectoine as a sole carbon source

To determine whether *V. diabolicus* can utilize ectoine as a sole carbon source, cells were grown at 37°C in M9 3% NaCl supplemented with 20 mM glucose, 20 mM ectoine or no carbon source, and growth was recorded for 30 h (**Fig. 5A**). *Vibrio diabolicus* grew in 20 mM ectoine as the sole carbon source with a 6 h lag phase and reached a final OD of 0.47 whereas growth in glucose had a lag phase <1 h and a final OD of 0.43 (**Fig. 5A**). Included in this analysis was growth in M9 3% NaCl without the addition of a carbon source as a negative control and as expected, no growth was observed (**Fig. 5A**). *Vibrio diabolicus* also grew in 20 mM ectoine as a sole carbon source at 30°C in M9 1% NaCl and M9 3% NaCl (data not shown). To demonstrate that the gene cluster was responsible for ectoine catabolism, an in-frame non-polar deletion of *V. diabolicus eutD* (OOKOHH_23780) was constructed. The *eutD* gene encodes the ectoine hydrolase, the first enzyme required for ectoine catabolism (**Fig. 1**). To show there was no overall growth defect in the Δ*eutD* mutant, this mutant was grown in M9 3% NaCl at 37°C with 20 mM glucose and showed an identical final OD to the wild-type strain (**Fig. 5B**). The *V. diabolicus* Δ*eutD* mutant was unable to grow in M9 3% NaCl with 20 mM ectoine at 37°C as the sole carbon source and confirmed that this gene is part of a gene cluster required for ectoine catabolism (**Fig. 5B**). In addition, *Vibrio natriegens*, a species that contained only *eutDE-asnC-ssd-atf*, showed no growth on M9 supplemented with 20 mM ectoine as the sole carbon source (data not shown).

**Fig. 5.**
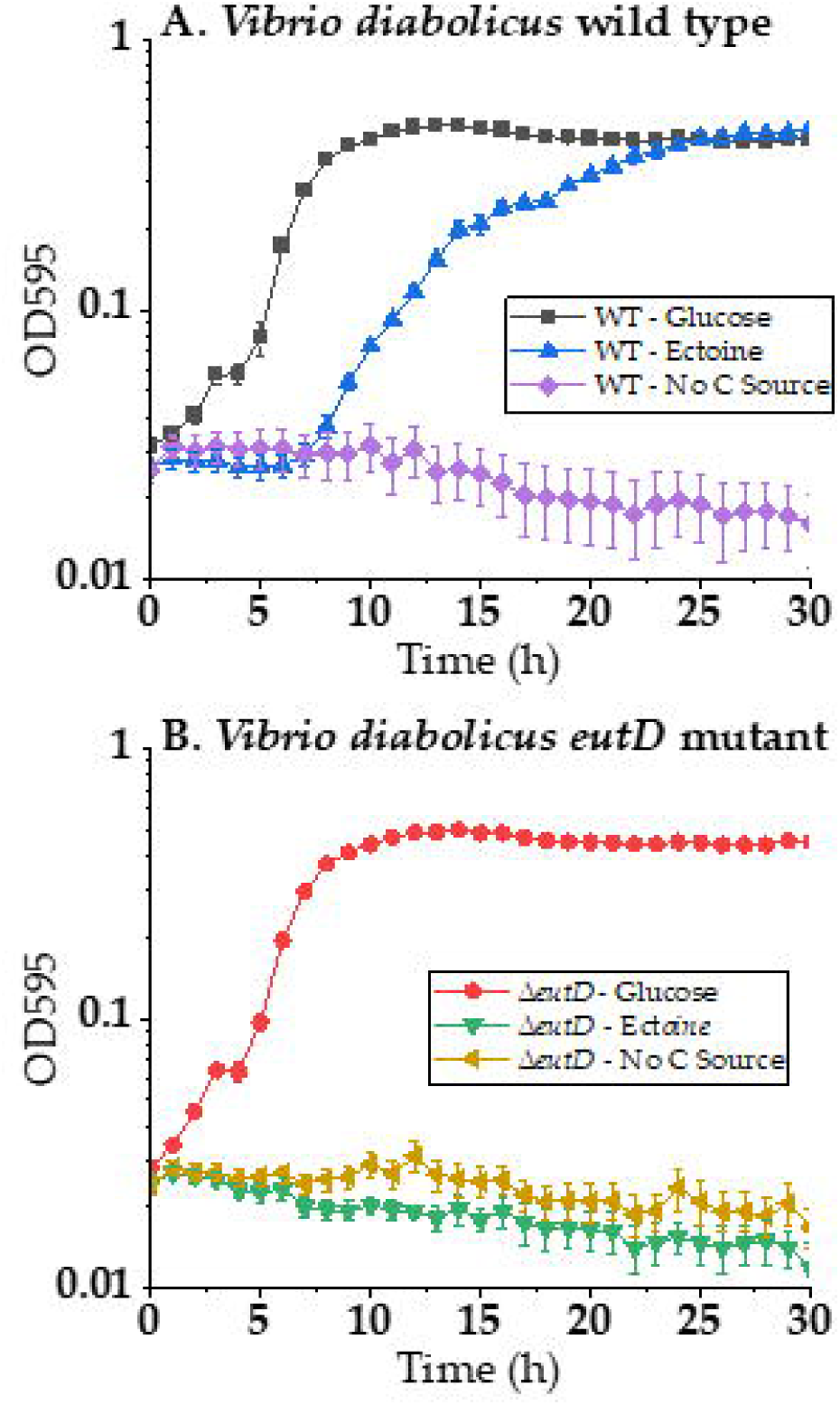
*Vibrio diabolicus* 3098 utilizes ectoine as a sole carbon source. A. Growth curve analysis of wild type in M9 3% NaCl at 37°C containing either 20 mM glucose (black line), 20 mM ectoine (blue line) or no carbon source (purple line). B. growth curve analysis of the *eutD* deletion mutant in M9 3% NaCl at 37°C containing either 20 mM glucose red line), 20 mM ectoine (green line) or no carbon source (yellow line).

Overall, our data show that *V. diabolicus* is both a producer and a consumer of ectoine, biosynthesizing it under salinity stress and capable of importing and catabolizing it as a nutrient source. These contradictory strategies co-exist because ectoine metabolism shifts between osmoprotection and nutrient acquisition based on environmental cues, including fluctuating salinity and nutrient availability. Distinct regulatory controls govern these processes, allowing intracellular ectoine pools to be preserved under salinity stress while enabling utilization of environmental ectoines as carbon and nitrogen sources when conditions permit. Although direct regulatory switching was not measured, the data are consistent with ectoine biosynthesis being induced by high NaCl concentration and catabolism was not affected by salinity, but consistent with substrate induction (controlled by regulator EnuR) similar to *R. pomeroyi*.

### Distribution of ectoine catabolism cluster among Proteobacteria, Thermodesulfobacteriota, Bacillota, and Actinomycetota

The EutD proteins from *V. diabolicus, R. pomeroyi,* and *S. meliloti* were used in BLAST searches of all available completed bacterial genomes in the NCBI RefSeq database as of June 2025. Within *Gammaproteobacteria,* there were 327 hits to genomic sequences with >60% amino acid identity among sequences; in *Betaproteobacteria*, there were 93 hits with >65% amino acid identity among sequences, and in *Alphaproteobacteria*, there were 580 hits with >55% amino acid identity among sequences annotated as ectoine hydrolase. Genomes that also contained EutE and EutB sequences were examined further using gene neighborhood analysis to identify additional genes involved in ectoine catabolism and transport. This was not an exhaustive search but one that concentrated on hits that contained both EutD, EutE, and EutB pathway genes, genomes with a species designation, and whole genome availability. A total of 575 EutD proteins from species representative of the diversity present within *Proteobacteria* (*Alphaproteobacteria, Betaproteobacteria,* and *Gammaproteobacteria*), and to a much lesser extent *Deltaproteobacteria, Thermodesulfobacteriota, Bacillota,* and *Actinomycetota* were examined further. An EutD homologue was present in 28 families of *Alphaproteobacteria (*species from saline, marine, and plant-associated environments), 4 families in *Betaproteobacteria* (species from rhizosphere soils and hydrophytes), and 5 *Gammaproteobacteria* families (mostly marine species from *Halomonadaceae* and *Vibrionaceae)*. In *Maridesulfovibrio bastinii* and *Oceanidesulfovibrio marinus*, two species of *Deltaproteobacteria (*renamed *Thermodesulfobacteriota),* a complete cluster was identified. The majority of these bacteria have never been experimentally shown to catabolize ectoine. In addition, we identified homologues of EutD in additional *Thermodesulfobacteriota* species, which contained 9-13 contiguous genes transcribed in the same direction usually with two *eutD* homologues, but lacked the *eutE* and *ssd* genes (**Fig. 6**). These putative ectoine catabolism clusters shared no synteny with the clusters described in *Proteobacteria* (**Fig 6**). Using the EutD and EutB proteins from this group in BLAST analyses, we identified putative ectoine catabolism gene clusters in *Bacillota* and *Actinomycetota* **(Fig. 7)**. Among species in the Order *Bacilliales*, the putative catabolism cluster was comprised of *pucR-eutBCD-atf* followed by *abgB-thrA-adh* and *eutD-msf*, all transcribed in the same direction (**Fig. 7**).

**Fig. 6.**
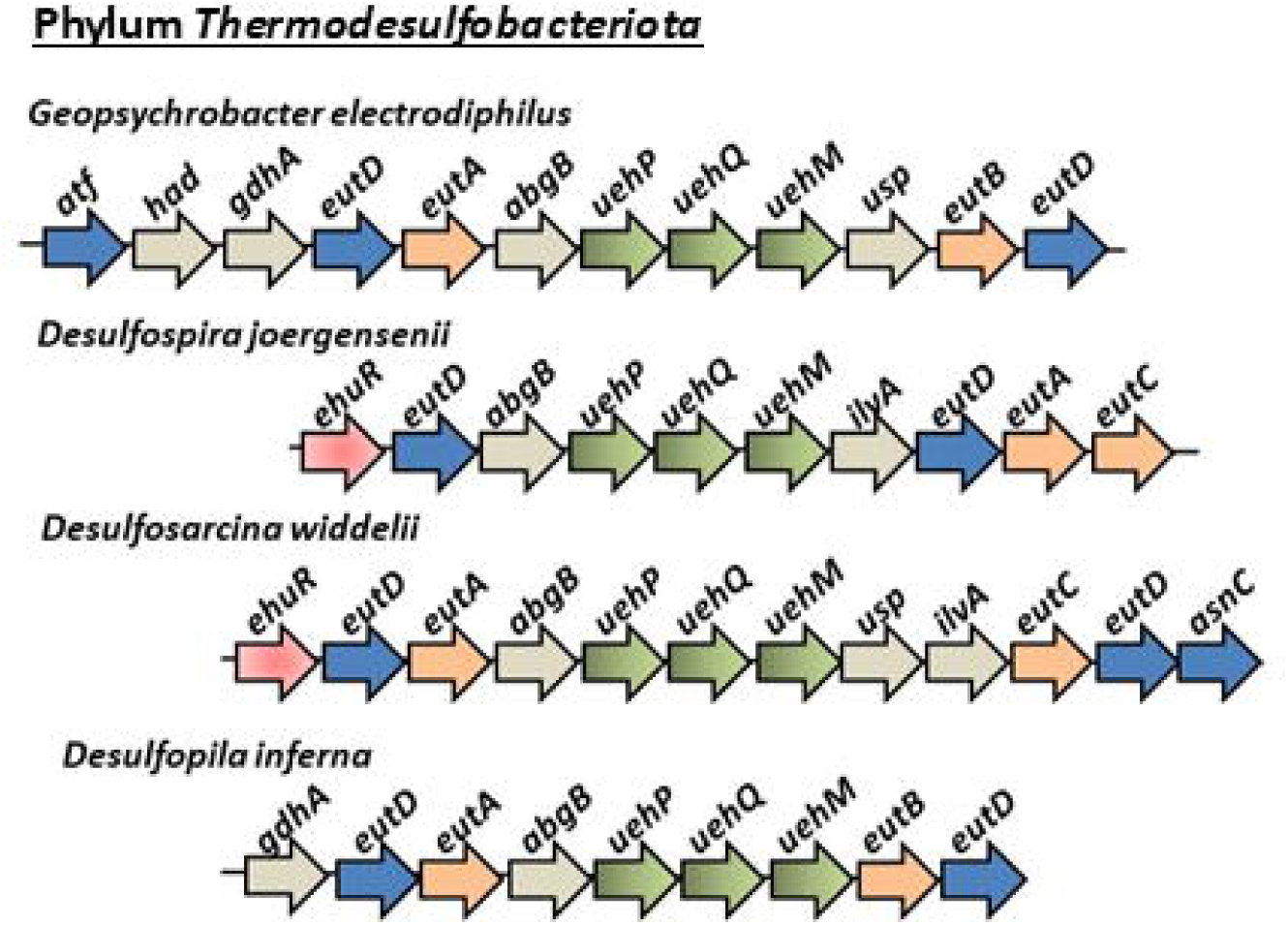
Ectoine and 5-hydroxyectoine catabolism gene clusters present in phylum *Thermodesulfobacteriota*. Analysis of *Thermodesulfobacteriota* putative ectoine catabolism clusters. Orange arrows represent *eutA, eutB, eutC*, and blue arrows represent *eutD, atf, and asnC*. Green arrows represent TRAP type transporter (*uehPQM*), red arrows indicate regulators (*enuR*), and grey proteins that have not been shown to be involved in ectoine catabolism.

**Fig. 7.**
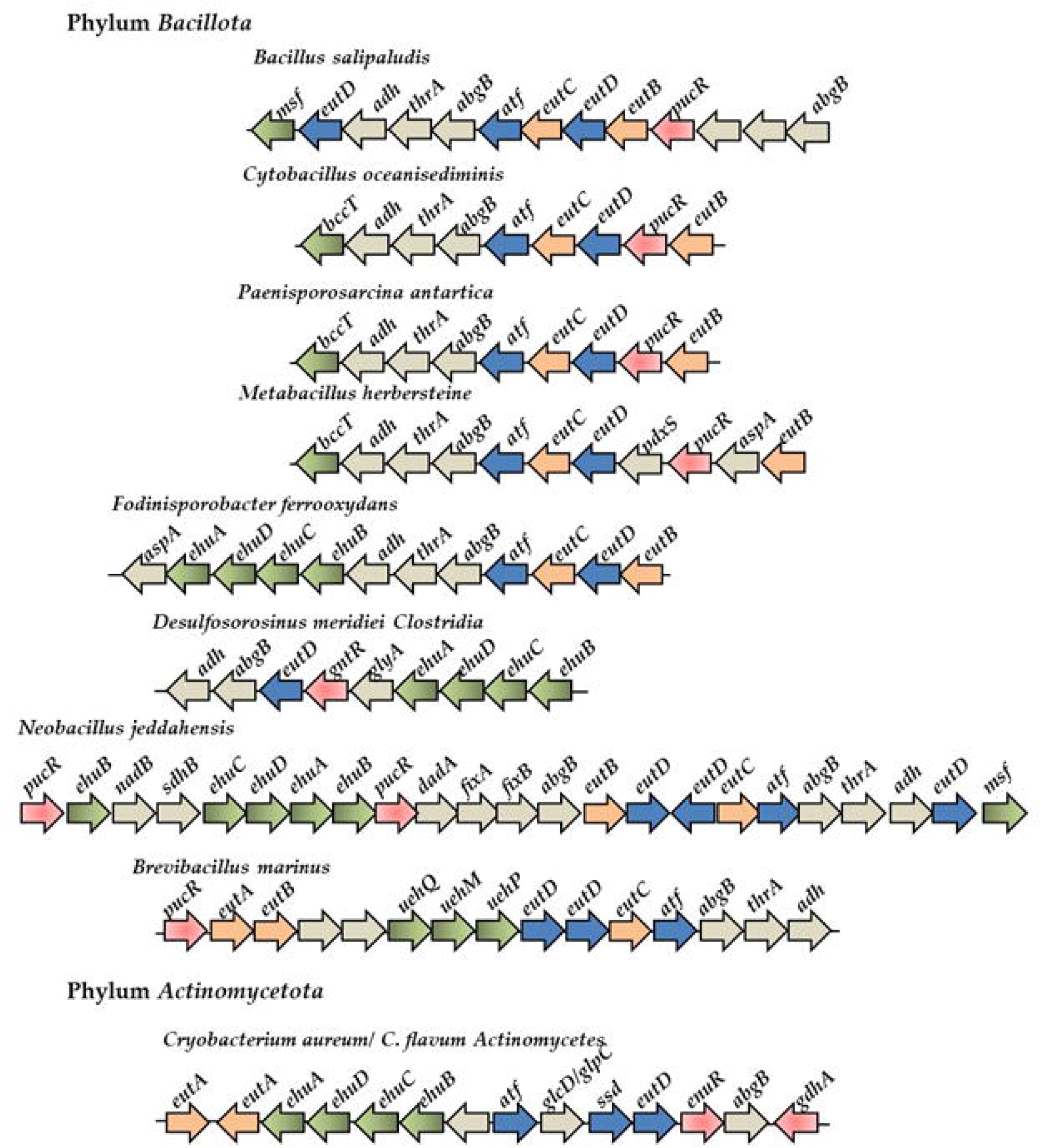
Ectoines catabolism gene clusters among Gram-positive bacteria. Orange arrows represent *eutB, eutC, and eutA* genes, blue arrows represent *eutD, atf,* and *ssd.* Green arrows transporter genes, red arrows indicate regulators and grey ORFs represents proteins whose functions in ectoine catabolism are unknown.

No EnuR homologue was present in *Bacillota*, but a PucR family transcriptional regulator clustered with the catabolism genes in most species examined (**Fig. 7**). PucR - type regulators were first described in *Bacillus subtilis* as a regulator of purine catabolism genes (52-54). They have not been shown to be involved in ectoine catabolism potentially indicating a possible novel function in *Bacillus*. The *bag* gene encodes a putative amidohydrolase potentially catalyzing hydrolysis of amide bonds, *thr* encodes an enzyme involved in reduction of aspartate beta semialdehyde to homoserine, and *adh* encodes an aldehyde dehydrogenase. These three genes were a feature of putative ectoine catabolism gene clusters identified in Gram-positive bacteria. Depending on the species, an MSF, BCCT, UehPQM, or EhuABCD was present within the clusters. Since these observations are based on sequence homology and genomic context, they remain tentative until physiological and biochemical validation.

### Evolutionary history of ectoine catabolism among bacteria

The 575 EutD sequences were aligned using CLUSTALW (55) and a phylogeny was constructed (**Fig. 8)**. For ease of reference, the major EutD groupings were labelled A to N on the tree and each group is described in detail with bootstrap values in subtrees in Figures S3-S9. The phylogeny showed that EutD sequences from each taxonomic class clustered together on the tree with notable exceptions (**Fig. 8 and labelled nodes in Fig. S2**). EutD from most *Pseudomonas* and *Stutzerimonas* species branched within the *Betaproteobacteria* clade (**Fig. 8**). EutD from *Gammaproteobacteria Marinobacterium* species had the most divergent EutD sequences among *Proteobacteria* and branched distantly from all other clades (**Fig. 8**). EutD from *Alphaproteobacteria* was present on divergent unrelated branches on the tree; clade C clustering with *Gammaproteobacteria* (A and B), clades D and E with *Betaproteobacteria* and *Gammaproteobacteria* (F and G), and clades H, I, J, K, and L forming the major *Alphaproteobacteria* grouping (**Fig. 8**). EutD proteins from *Thermosulfobacteriota* and *Bacillota* clustered together on divergent branches distantly from *Proteobacteria* (**Fig. 8**). Nested within this latter grouping is EutD from two genera formally designated *Deltaproteobacteria,* which indicates that the region was acquired from *Alphaproteobacteria* (**Fig. 8**).

**Fig. 8.**
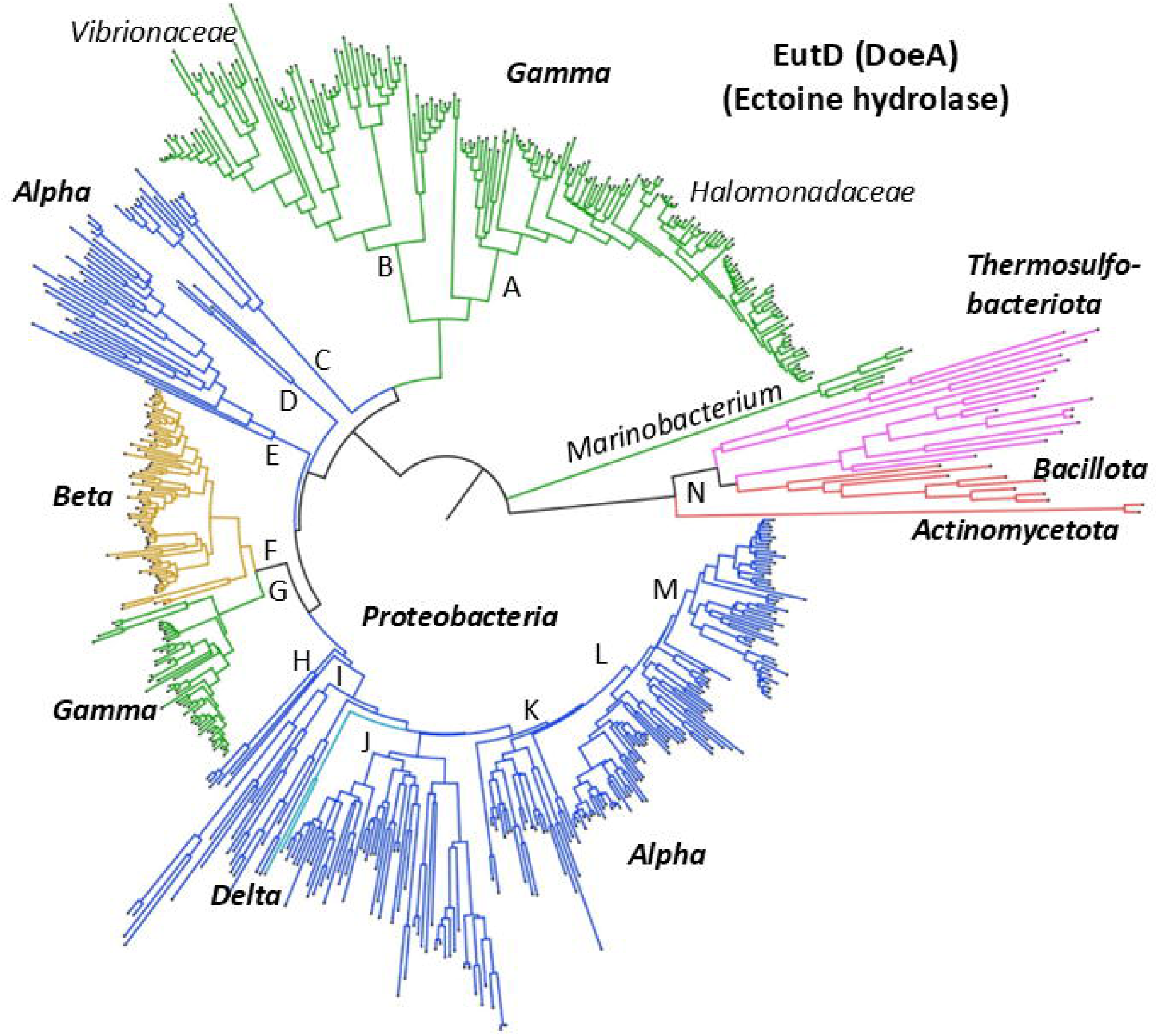
Phylogeny of EutD (DoeA) from 575 taxa. The evolutionary history was inferred using the Neighbor-Joining method with distances computed using the JTT matrix-based method. The optimal tree with the sum of branch length = 38.256 is shown. The tree was drawn to scale, with branch lengths in the same units as those of the evolutionary distances used to infer the phylogenetic tree. The rate variation among sites was modelled with a gamma distribution (shape parameter = 3.00). The complete deletion option was applied resulting in a final data set comprising 378 positions. Evolutionary analyses were conducted in MEGA12 utilizing up to 3 parallel computing threads. The tree with each node labelled can be found in Fig. S2 and in subtrees Fig. S3-S9.

EutD was present in several *Vibrionaceae* genera and clustered along genus designation with a cluster present in *Vibrio, Photobacterium, Grimontia, Enterovibrio*, and *Salinovibrio* on chromosome 2 (**Fig. S3**). EutD from *Photobacterium atrarenae* nested within *Vibrio* EutD proteins, indicating the genes were acquired by horizontal transfer. Only a limited number of *Vibrio* species contained the entire cluster of catabolism, transport, and regulatory genes: *V. diabolicus*, *V. antiquarius, V. chemagurensis, V. hepatarius*, and *V. proteolyticus* **(Fig. S3).** Both *V. diabolicus* and *V. antiquarius* contain strains isolated from deep-sea hydrothermal vents, and are synonym, with all sequenced genomes containing the catabolism cluster (49). *Vibrio diabolicus*, *V. antiquarius,* and *V. chemagurensis* are closely related to *V. alginolyticus* in the Harveyi clade. Among all *V. alginolyticus* strains (>1,200 genomes), a six-gene cluster *(enuR*-*eutDE-asnC-ssd-atf)* is present at the same genome location as in *V. diabolicus* on chromosome 2 (**Fig. S3**). In *Vibrio* species that contained only the *eutD_asnC-ssd-atf* cluster, this was present on chromosome 1 (**Fig. S3**). In species of *Photobacterium, Grimontia, Enterovibrio and Salinovibrio*, the *enuR*-*eutDE-asnC-ssd-atf* cluster was present on chromosome 2. These data indicate that the cluster has a complex history that involves gene loss and horizontal and vertical transmission. The EutD phylogeny showed that EutD from *Vibrionaceae* shared a most recent common ancestor with *Aeromonadaceae*, but in this group, the TRAP transporter genes were in an operon (**Fig. S3**). EutD from 9 *Pseudomonas* species branched with EutD from *Nitrincola alkalilacustris* divergent from vibrios and contained a 12-gene cluster like *V. diabolicus* (**Fig. S3**). Group A from Figure 8 was comprised of EutD from all members of *Halomonadaceae*, with most EutD proteins in this group clustered along taxonomic lines (**Fig S4**). Among *Halomonadaceae*, an *eutA* homologue was absent and transporter genes typically did not cluster with the catabolism genes (**Fig. S4**). In addition, *eutBC* was absent from all *Vreelandella* species and a third of the *Halomonas* species suggesting gene loss in these genera. In our analysis, the *Halomonas elongata teaABC* cluster showed limited distribution within *Halomonadaceae*, whereas the *uehPQM* and *ehuABCD* transporters were more prevalent. EutD from 12 *Salinicola* species formed two related branches, one branch contained four species with UehPQM and the other branch eight species with EhuABCD (F**ig. S4**). These data suggest that the catabolism cluster is ancestral, and the transporter genes were acquired separately. EutD sequences from *Roseovarius* and *Nitratireductor* species formed a divergent clade (clade C) from clades A and B; these *Alphaproteobacteria* contained the *uehPQM* and *ehuBCDA* operons, respectively (**Fig. S3**).

Among *Betaproteobacteria*, EutD was present in in *Burkholderiaceae (Burkholderia, Caballeronia, Paraburkholderia, Trinickia),* one species of *Comamonadaceae (Verminephobacter eiseniae),* and four species of *Rhodocyclaceae* and *Zoogloeaceae* (F**ig. S5**). *Burkholderiaceae* contain species that are highly versatile occupying many ecological niches in terrestrial and aquatic environments. *Burkholderia* contain pathogenic species as well as insect gut symbionts, and *Caballeronia* and *Paraburkholderia* consist of beneficial plant-associated symbiotic species (56-59). EutD from *Burkholderia* species all clustered on the same branch but no ectoine transporter was identified. Whereas in *Paraburkholderia, Caballeronia,* and *Trinickia*, all of which contained *ehuBCDA* divergently transcribed from the catabolism genes, no *eutA* homologue was present (F**ig. S5**). In a subset of *Paraburkholderia* species, an *ehuABCD* cluster was present and this cluster also encoded an *eutA* homologue (**Fig. S5**). Nested within EutD from *Burkholderiaceae* was EutD from *Verminephobacter eiseniae* that contained a catabolism cluster identical to *Paraburkholderia,* indicating horizontal transfer (F**ig. S5**). EutD from *Aromatoleum* and *Denitromonas,* clustered together on two divergent branches, and the catabolism clusters contained the UehPQM transporter. EutD from most *Pseudomonas* and *Stutzerimonas* species shared a most recent common ancestor with EutD from *Betaproteobacteria,* with conserved synteny and the presence of the EhuBCDA transporter **(Fig. S5)**. Overall, the data suggest that ectoine catabolism is ancestral to *Burkholderiaceae* and the transporter genes acquired separately whereas in *Pseudomonas* species, the region was acquired twice.

EutD was the most phylogenetically widespread among *Alphaproteobacteria* present in at least 28 diverse families. The phylogeny showed that EutD from *Alphaproteobacteria* has an ancient and varied history, forming highly divergent unrelated lineages (**Fig. 8**). Gene content and order were highly conserved within many lineages (**Fig. S6–S8**) but differed from those in *Gammaproteobacteria* and *Betaproteobacteria*; for example, gene order was *eutABC* rather than *eutBCA* as observed in *Gammaproteobacteria*. Four different transporter clusters (*uehPQM, uehP, uehQM, ehuABCD,* and *ehuBCDA)* were present within the catabolism genes depending on the lineage. EutD clusters in H, I and J contained species with *uehPQM*; clusters K and L contained *ehuABCD,* whereas cluster M contained *ehuBCDA* (**Fig. S6, S8**). The *uehP, uehQM* genes were present in *Thalassospira* species (**Fig. S7**). Since the transporter genes in most cases were embedded within the catabolism genes, the most parsimonious evolutionary explanation is that the catabolism and transporter genes were acquired together (**Fig. S8**). The phylogeny distribution and gene cluster analysis suggest convergent evolution within *Alphaproteobacteria* with long term vertical transmission and retention. EutD sequences from the genus *Paracoccus* formed two divergent branches. Species containing EhuABCD clustered with other members of *Paracoccaceae* (**Fig. S8 group K**), whereas those containing UehPQM grouped with *Roseobacteraceae* (**Fig. S7 group J**). EutD from *Verminephobacter aporrectodeae*, a *Betaproteobacteria,* clustered within the EutD *Alphaproteobacteria* group L, indicating horizontal acquisition in this species (**Fig. S8**).

Ectoine catabolism was not previously described in Gram-positive bacteria. Hence, the discovery of putative ectoine catabolism clusters within *Bacillota* and *Actinomycetota* genomes was unexpected (**Fig. 7**). From the small group of species that we examined in the *Bacillaceae*, the operon *eutD-eutC-atf* is conserved, followed by the operon *abgB-thrA-adh,* which was present in most species examined (**Fig. 7**). In these species, an *eutE* homologue is absent from the genomes but *eutA, eutB* and an additional *eutD* homologue are often contiguous with the other pathway genes (**Fig. 7**). A BCCT, EhuABCD, or UehPQM transporter also clustered with the putative ectoine catabolism genes depending on the species (**Fig. 7**). The EutD proteins from Firmicutes formed the deepest branches within the tree with long branch lengths, suggesting an evolutionary origin unrelated to *Proteobacteria* (**Fig. 8 and Fig. S9**). This is also evidenced by the lack of synteny between Gram-negative and Gram-positive bacteria and the absence of an EnuR homologue among *Bacillota* and the presence of the PucR regulator a known regulator of purine catabolism in *Bacillus subtilis* (**Fig. 7**) **(52-54)**. The long branch lengths reflect gaps in current sampling and should be resolved as additional Gram-positive genomes are sequenced, providing a clearer picture of ectoine catabolism in this group.

The identification of putative ectoine catabolism genes among Gram-positive bacteria prompted us to investigate whether members of the Domain *Archaea* also harbored putative ectoine catabolism clusters. We used EutD (DoeA) (WP_133333823.1) from *Bacillus salipaludis* in BLAST analysis of *Archaea*, which returned over a hundred hits with amino acid identity > 55% and >97% query coverage (**Fig. S10**). The putative EutD proteins were within clusters of genes that included aminopeptidase and aminotransferase genes as well as a BCCT transporter with the clusters conserved between genera (**Fig. S11**).

## Conclusions

In summary, this study demonstrated that *V. diabolicus* is a producer of ectoine under osmotic stress and a consumer of ectoine under nutrient-limiting conditions. This metabolic flexibility is an important evolutionary strategy for this species. The conservation of ectoine catabolism and regulatory clusters strongly suggests that many species are capable of catabolizing ectoine. Phylogenetic analysis revealed that EutD has an ancient, highly dynamic evolutionary history shaped by lineage-specific diversification, convergent evolution, and infrequent horizontal gene transfer. Within *Gammaproteobacteria,* distinct patterns of both vertical and horizontal transfer are evident as well as gene loss: EutD is present in a limited number of *Vibrio* species but in the majority of *Halomonadaceae*, with EutD from *Pseudomonas* showing evolutionary origins within *Gammaproteobacteria* and *Betaproteobacteria*. Ectoine catabolism evolution among *Alphaproteobacteria* displayed the greatest distribution and diversity forming multiple deep, unrelated lineages, which suggests multiple acquisitions followed by long-term retention. Different transporter systems were embedded within catabolic gene clusters in *Alphaproteobacteria* suggesting co-acquisition and convergent evolution in this group. The diversity of transporters associated with different taxonomic groups is consistent with modular evolution of ectoine utilization with a conserved catabolic core paired with lineage-specific transporters allowing for adaptation based on ectoine availability/concentration. Finally, although genomic and phylogenetic analyses support widespread ectoine catabolism among Bacteria and Archaea, experimental validation is currently limited to a small number of *Proteobacteria* taxa. Putative ectoine catabolism clusters identified in Gram-positive bacteria and *Archaea* lack any direct biochemical confirmation and may represent independently evolved or functionally divergent pathways. Accordingly, these putative ectoine clusters are considered provisional until experimentally evaluated.

## MATERIALS AND METHODS

### Bacterial strains

*Vibrio diabolicus* strain 3098 was used in this study (18). Growth media used was either Luria-Bertani medium (LB) (Fisher Scientific, Fair Lawn, NJ) or M9 minimal medium (47.8 mM Na_2_HPO_4_, 22 mM KH_2_PO_4_, 18.7 mM NH_4_Cl, 8.6 mM NaCl; Sigma Aldrich) supplemented with 2 mM MgSO_4_, 0.1 mM CaCl_2_, and 20 mM glucose as the sole carbon source (M9G). Media had a final NaCl concentrations of 3% (wt/vol) NaCl or 2% (wt/vol) NaCl for *V. diabolicus* 3098 and *V. natriegens* ATCC 14048 respectively, unless stated otherwise. *Escherichia coli* strains were grown in LB supplemented with 1% (wt/vol) NaCl (LB 1%) at 37°C. To generate a deletion mutant, the diaminopimelic acid (DAP) auxotroph, *E. coli* β2155 *λpir* was used (60). When grown, this strain was supplemented with 0.3 mM DAP. When specified, chloramphenicol (Cm) was added to media at a concentration of 12.5 µg/mL.

### Construction of deletion mutant

An in-frame nonpolar deletion mutant of *eutD* was generated in *V. diabolicus* 3098. The mutant was constructed using SOE PCR and allelic exchange (61) using the primers AB fwd_*eutD (*gtggaattcccgggagagctATATGCATGACTGACGGTTTG) and AB rev_*eutD* (ataagtggcgATGCAAAGCAACGCCTCTC) and CD fwd_*eutD* (tgctttgcatCGCCACTTATTTGTTAAACAC) and CD rev_*eutD* (accgcatgcgatatcgagctAAAATCGCACATGGGTACTAAG) to construct a truncated in-frame deletion in the *eutD* gene (45-bp of the 1188-bp gene). Using the Gibson assembly protocol, the truncated gene product was ligated with the suicide vector pDS132 (pDSΔ*eutD*) and transformed into *E. coli* β2155 *λpir* (DAP auxotroph) as described in (62). *Escherichia coli* β2155 *λpir* pDSΔ*eutD* was conjugated into *V. diabolicus* using a contact-dependent biparental mating by cross-streaking the two strains on LB agar with 1% NaCl and 0.3 mM DAP. The plasmid vector undergoes homologous recombination into the *V. diabolicus* genome, since it does not contain the *pir* gene required for replication of the pDSΔ*eutD* plasmid. Following a series of selections on chloramphenicol and sucrose plates, the recombinant clones that underwent double crossover (Δ*eutD*) were selected for the phenotype SacB^r^Cm^s^. An in-frame deletion of *eutD* was confirmed by colony PCR and DNA sequencing.

### Preparation of cellular extracts and proton nuclear magnetic resonance (^1^H-NMR)

Wild-type *V. diabolicus* 3098 was grown overnight at 37°C in M9G 1% NaCl, M9G 5% NaCl, or M9G 5% NaCl + 1 mM choline. Stationary phase cells were pelleted and washed twice with PBS. To ensure lysis of cells, three freeze-thaw cycles were completed with cell pellets. Cell pellets were then resuspended in 750 µL ethanol and debris was pelleted by centrifugation. The ethanol solutions were transferred to clean tubes and evaporated under vacuum. To resuspend the pellet, 700 µL deuterium oxide (D_2_O) was used to vortex the pellet, and insoluble material was removed by centrifugation for 10 min at 12,000 rpm. Extracts were examined on a Bruker AVANCE 600NMR spectrometer with 16 scans per spectrum. Data were analyzed using MestReNova (Mnova) NMR software.

### Osmotic stress growth analysis

*Vibrio diabolicus* was grown to stationary phase in M9G 1% NaCl at 37°C. Fresh M9G 1% NaCl media was inoculated (2%) from the stationary culture and allowed to grow to OD ∼ 0.55 at 30 or 37°C with aeration by shaking. The exponential cultures were inoculated 1:40 into 200 µL fresh M9G media containing 0 to 7% NaCl. The compatible solutes, glycine betaine (GB), dimethylsulfoniopropionate (DMSP), and ectoine were added to M9G 7% NaCl growth media at a final concentration of 1 mM. Plates were incubated at 30 or 37°C with hourly shaking for 1 min. The optical densities were measured at 595 nm hourly for 24 h using a Tecan Sunrise microplate reader and Magellan plate reader software (Tecan Systems, Inc., San Jose, CA).

### Growth on ectoine as sole carbon source

For growth analysis, *V. diabolicus* and *V. natriegens* were grown overnight in minimal media (M9) in 3% NaCl and 20 mM glucose at 37°C. These cultures were then diluted 1:50 into M9 and grown to OD of ∼0.5. These cells were then washed twice with PBS to remove glucose. Washed cultures were inoculated 1:40 into 200 µL M9 media supplemented with 3% NaCl, and specified carbon source (20 mM glucose or 20 mM ectoine). Plates were incubated at 37°C with intermittent shaking for 1 min every hour. The optical densities were measured at 595 nm hourly for a total of 30 h using a Tecan Sunrise microplate reader and Magellan plate reader software (Tecan Systems Inc., San Jose, CA).

### Phylogenetic analysis

To determine the distribution and evolutionary history of the ectoine catabolism cluster among bacteria, the EutD protein from *V. diabolicus* 3098, *R. pomeroyi* and *S. meliloti* were used in BLAST searches of the NCBI bacterial genome database June 2025. Homologues of EutD were identified within *Gammaproteobacteria,* then *Betaproteobacteria* and *Alphaproteobacteria*. The most divergent EutD proteins identified in each subphylum were used in reciprocal searches of the RefSeq select protein database. Among EutD proteins identified in each species were further examined using EFI-Gene neighborhood tool (GNT), a program that performs sequence similarity cluster analyses (63). Only EutD proteins that clustered with additional ectoine catabolism genes and were within strains identified to the species level were examined further. Not all EutD from certain genera were included as hundreds of closely related sequences were available, as in *Halomonas, Pseudomonas, Paracoccus, Sinorhizobium, Rhizobium, Mesorhizobium,* and *Agrobacterium*. In addition, the analysis of possible ectoine catabolism gene clusters among Gram-positive bacteria was not comprehensive. Therefore, the analysis of the distribution of EutD is not an exhaustive representation of possible ectoine catabolism pathways among bacteria. EutD sequence alignments were completed using CLUSTAL (55) with the program MEGA 12 (64). The evolutionary history of proteins in this study were inferred using the Neighbor-Joining method (65) and evolutionary distances were computed using the JTT matrix-based method. Ambiguous positions were removed for each sequence pair (complete deletion option).

## Data Availability

All datasets are available upon request.

## ACKNOWLEDGMENTS

K.E.B.L. was funded in part by the Chemistry-Biology Interface Predoctoral Training Program (grant 5T32GM008550).

