## Supplementary tables for "Deciphering the evolutionary history of ectoine catabolism, a compatible solute utilized by *Vibrio diabolicus* as an osmoprotectant and a nutrient source"

**Table S1.** Ectoine and hydroxyectoine catabolism proteins identified in *Vibrio diabolicus* with homology to *Halomonas elongata* and *Ruegeria pomeroyi.* Names in brackets are nomenclature from *Halomonas elongata*

| ***Vibrio diabolicus*** | **Protein ID** | **Product** | ***Halomonas elongata*** | ***Rugeria pomeroyi*** |
|---|---|---|---|---|
| **EutA** | WP_134645445.1 | Asp/Glu racemase | - | 31% |
| **EutC** | WP_256936410.1 | Cyclodeaminase | 57% | 46% |
| **EutB** | WP_006741390.1 | Hydroxyectoine dehydratase | 57% | 50% |
| **Atf (DoeD)** | WP_074190436.1 | Aspartate aminotransferase | 68% | 60% |
| **Ssd (DoeC)** | WP_243328035.1 | NAD-dependent succinate-semialdehyde | 63% | 52% |
| **AsnC (DoeX)** | WP_006741393.1 | AsnC family regulator | 66% | 52% |
| **UehP** | WP_104972313.1 | TRAP substrate-binding protein | 41% **(TeaA**) | 42% **(UehA)** |
| **EutE (DoeB)** | WP_006741395.1 | diaminobutanoate deacetylase | 64% | 55% |
| **EutD (DoeA)** | WP_005392731.1 | Ectoine hydrolase | 63% | 62% |
| **EnuR** | WP_258624098.1 | MocR family regulator with PLP-dependent domain | 41% | 34% |
| **UehQ** | WP_145472721.1 | TRAP transporter small permease | 38% (**TeaB**) | 44% **(UehB)** |
| **UehM** | WP_005392736.1 | TRAP transporter large permease | 64% (**TeaC**) | 67% **(UehC)** |

**Table S2.** *Vibrio diabolicus* NaCl tolerance range and growth at different temperatures. Final OD values and maximum growth rates are displayed.

| M9 30°C | OD | Growth rate |
|---|---|---|
| 0 | 0.14 | 0.14 |
| 1%NaCl | 0.56 | 0.57 |
| 2%NaCl | 0.64 | 0.50 |
| 3%NaCl | 0.63 | 0.60 |
| 4%NaCl | 0.64 | 0.41 |
| 5%NaCl | 0.66 | 0.47 |
| 6%NaCl | 0.56 | 0.32 |
| 7%NaCl | 0 | 0 |

| M9 37°C | OD | Growth rate |
|---|---|---|
| 0 | 0 | 0 |
| 1%NaCl | 0.49 | 0.51 |
| 2%NaCl | 0.70 | 0.69 |
| 3%NaCl | 0.72 | 0.90 |
| 4%NaCl | 0.83 | 0.83 |
| 5%NaCl | 0.67 | 0.58 |
| 6%NaCl | 0.76 | 0.39 |
| 7%NaCl | 0 | 0 |
