## supplementary figures for "Deciphering the evolutionary history of ectoine catabolism, a compatible solute utilized by *Vibrio diabolicus* as an osmoprotectant and a nutrient source"

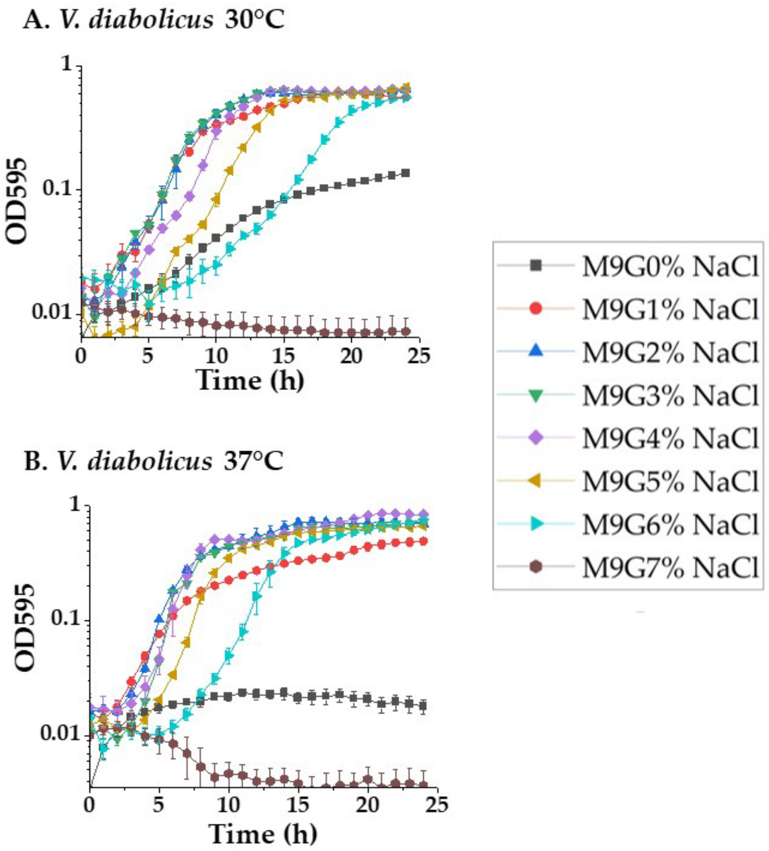


**Fig. S1.** ***Vibrio diabolicus*** **3098 NaCl tolerance range and growth at different temperatures.** Growth curves of *V. diabolicus* strain 3098 in minimal media (M9) with glucose (M9G) with a final concentration of 0% to 7% NaCl. Growth was measured every hour for 24 h at **A.** 30°C and **B.** 37°C.

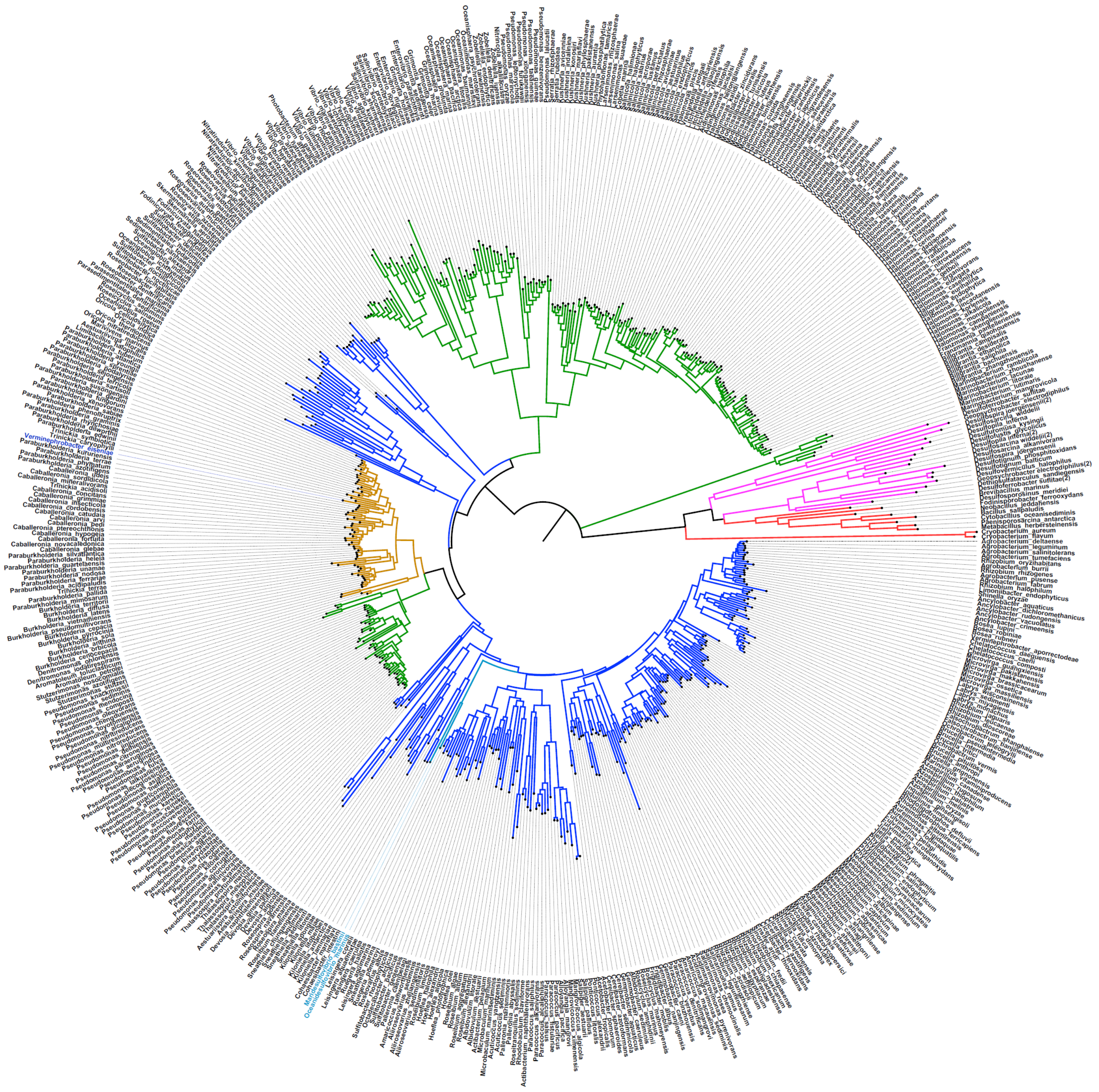

**Fig. S2 Labelled Fig. 8. Phylogeny of EutD (DoeA) from 575 taxa.** The evolutionary history was inferred using the Neighbor-Joining method with distances computed using the JTT matrix-based method. The complete deletion option was applied resulting in a final data set comprising 378 positions. Evolutionary analyses were conducted in MEGA12 [4] utilizing up to 3 parallel computing threads. Branch coloring is as follows: Green=Gammaproteobacteria, Yellow=*Betaproteobacteria*, Blue=*Alphaproteobacteria*, Aqua=*Deltaproteobacteria*, Red= *Bacillota* and *Actinomycetota,* Mauve=*Thermosulfobacteriota*

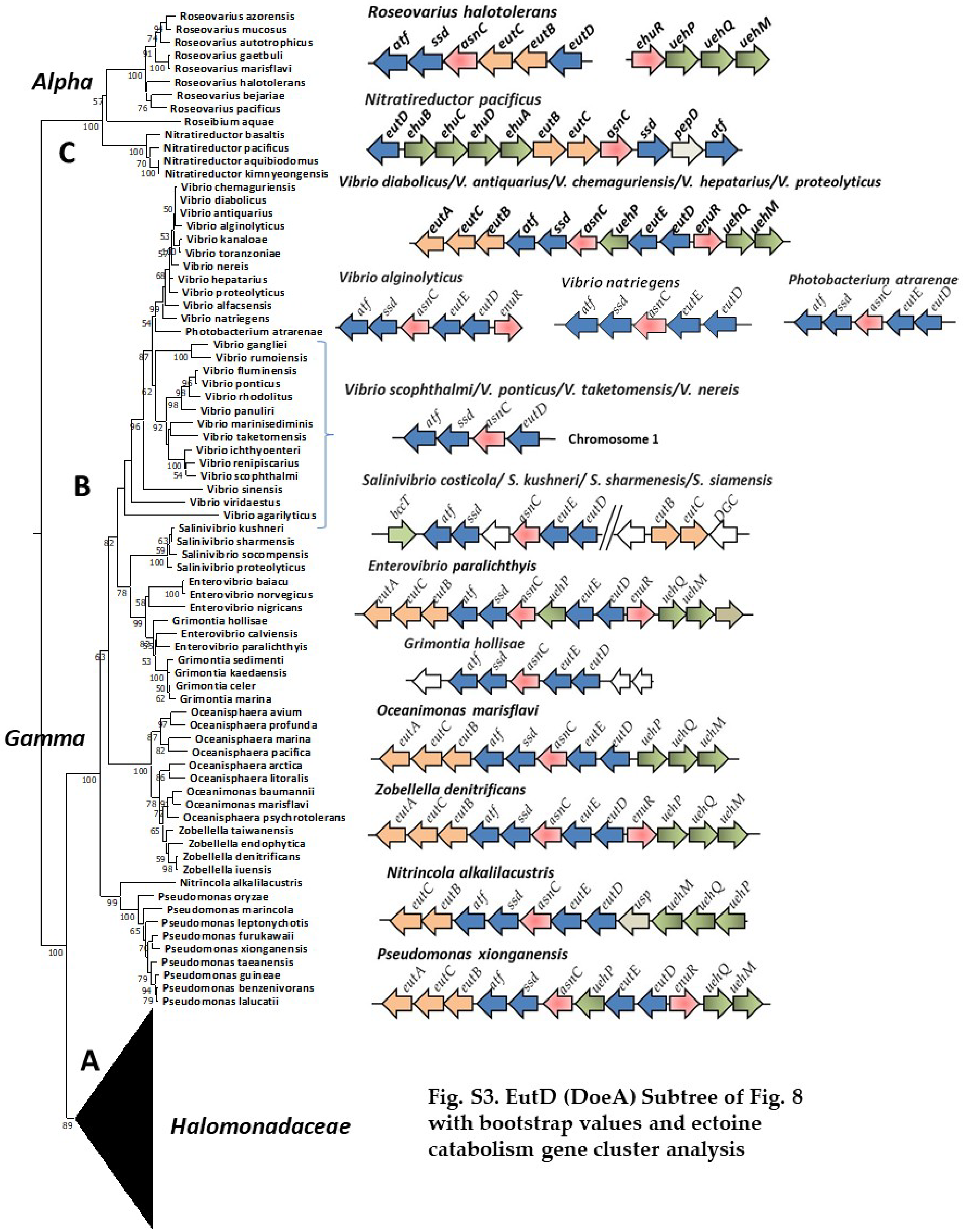


**Fig. S3. EutD (DoeA) Subtree of Fig. 8 with bootstrap values and ectoine catabolism gene cluster analysis**

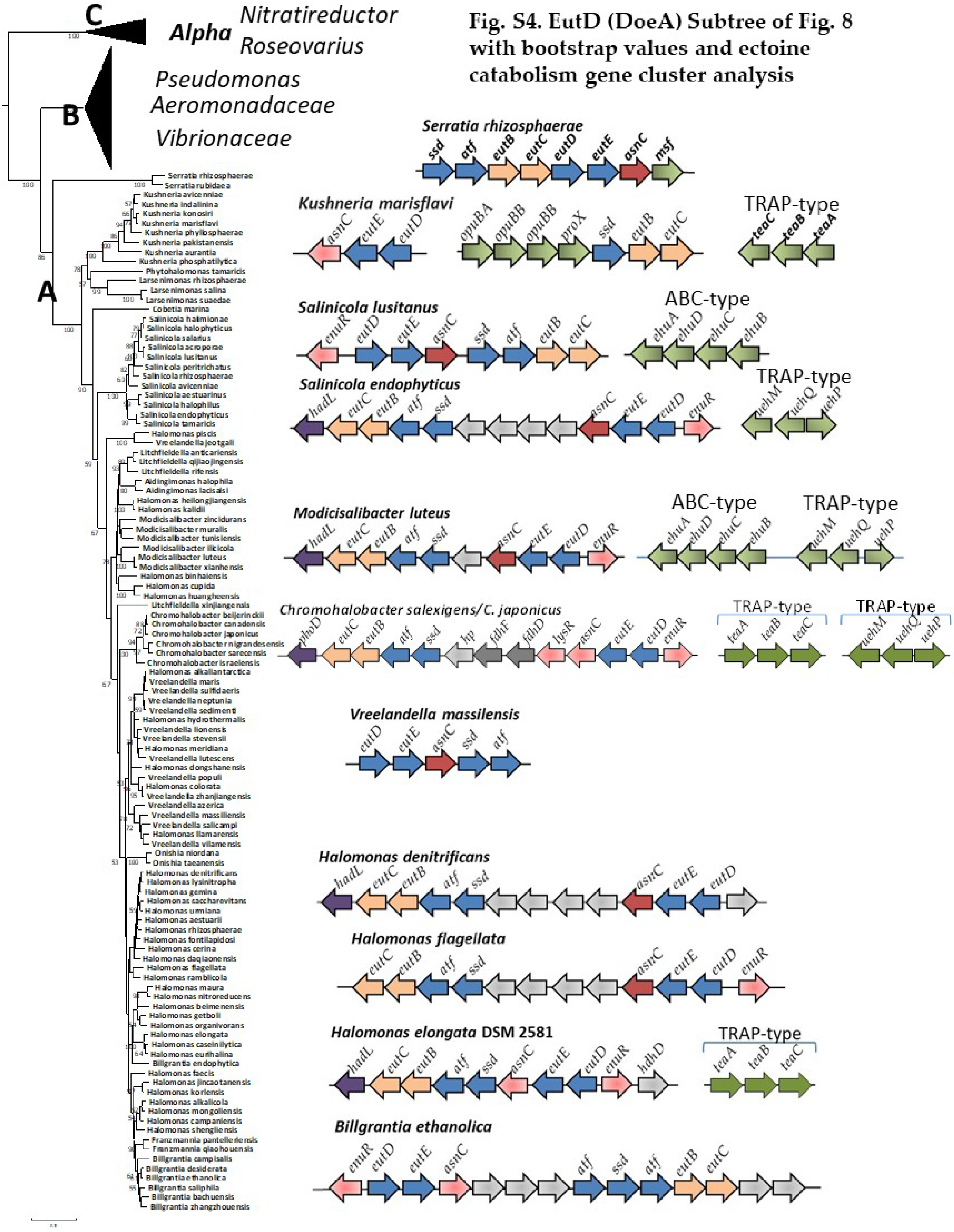


**Fig. S4. EutD (DoeA) Subtree of Fig. 8 with bootstrap values and ectoine catabolism gene cluster analysis**

Fig. S5. EutD (DoeA) Subtree of Fig. 8 with bootstrap values and ectoine catabolism gene cluster analysis

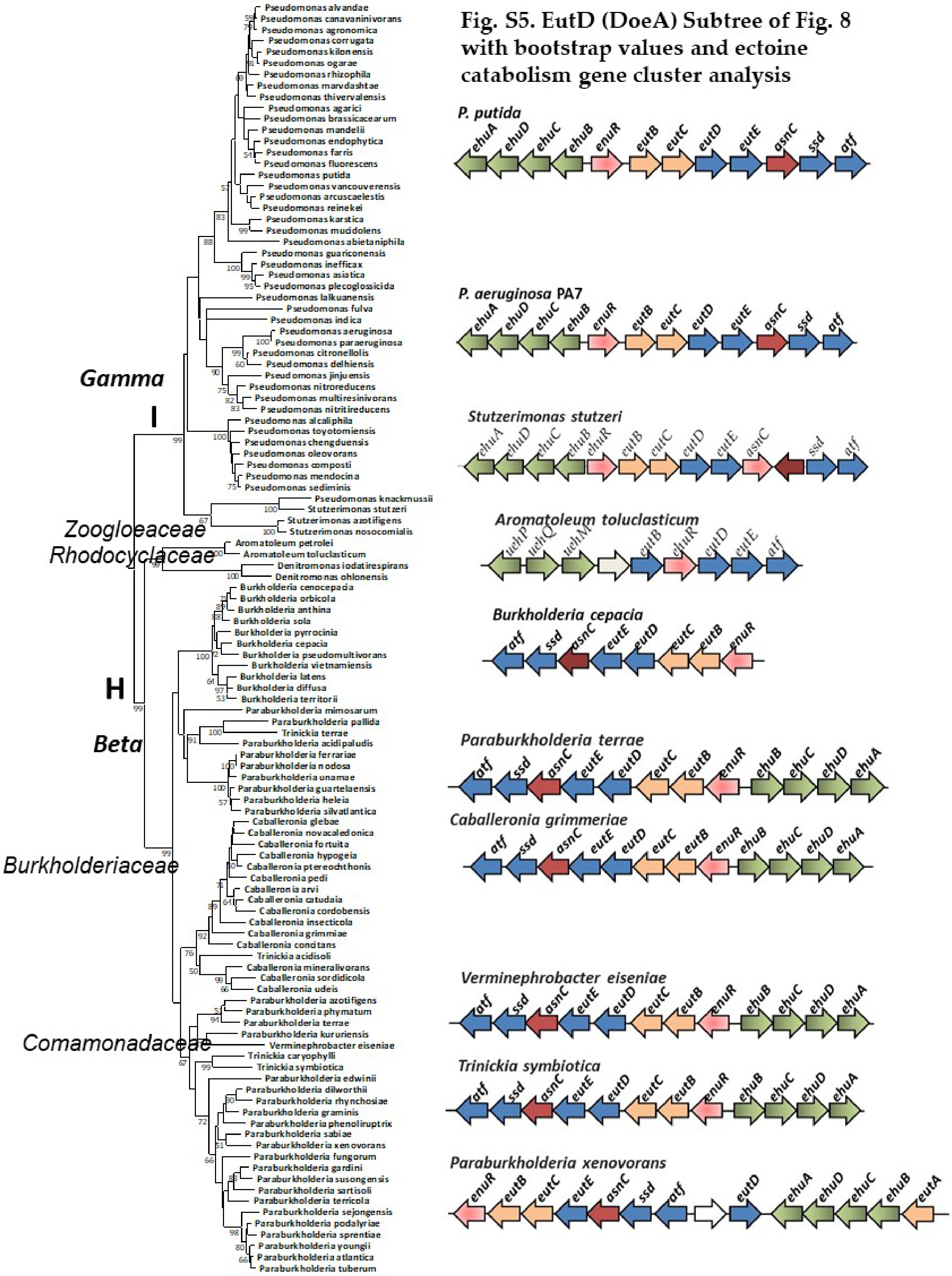

**Fig. S6. EutD (DoeA) Subtree of Fig. 8 with bootstrap values and ectoine catabolism gene cluster analysis**

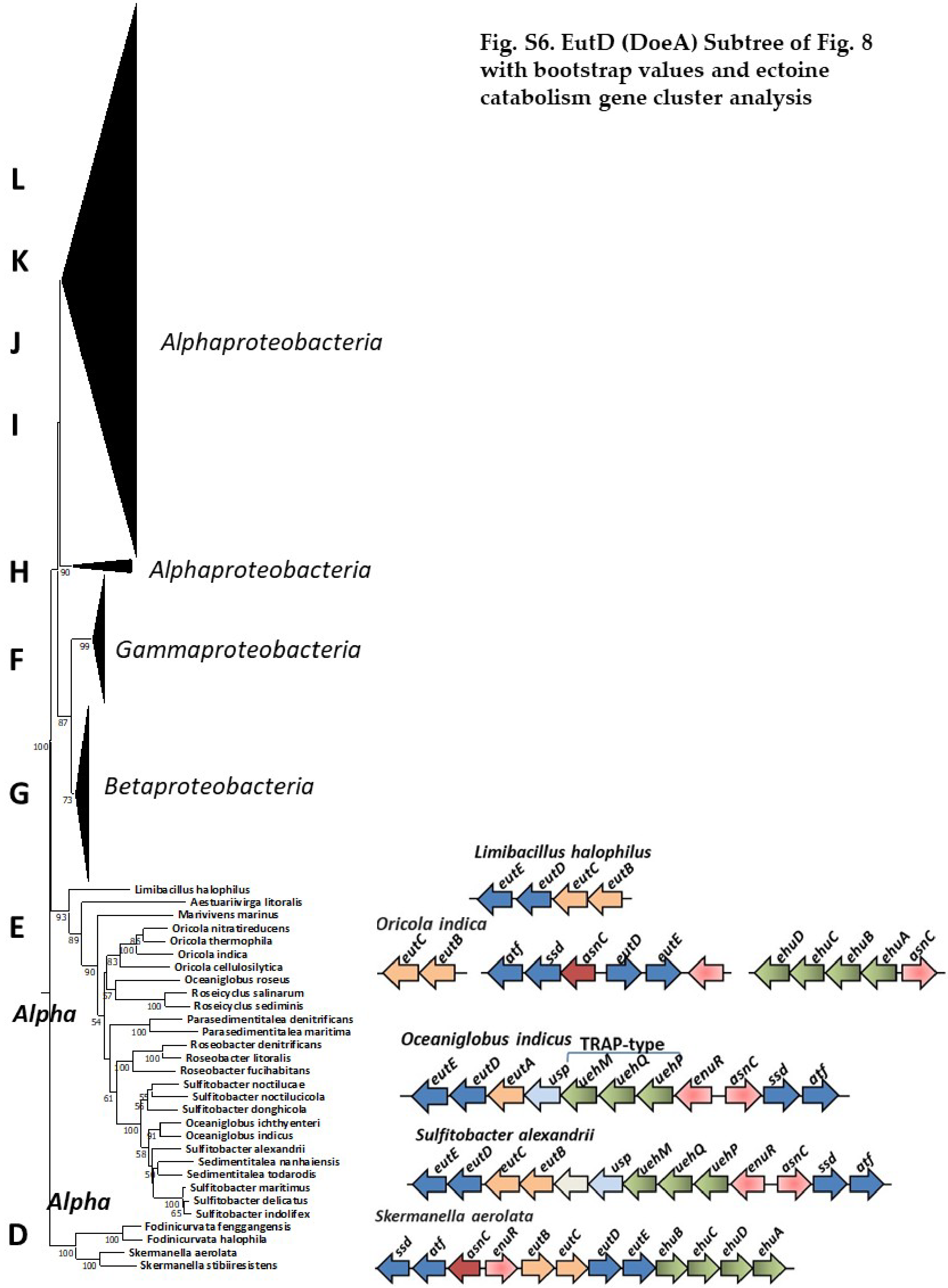

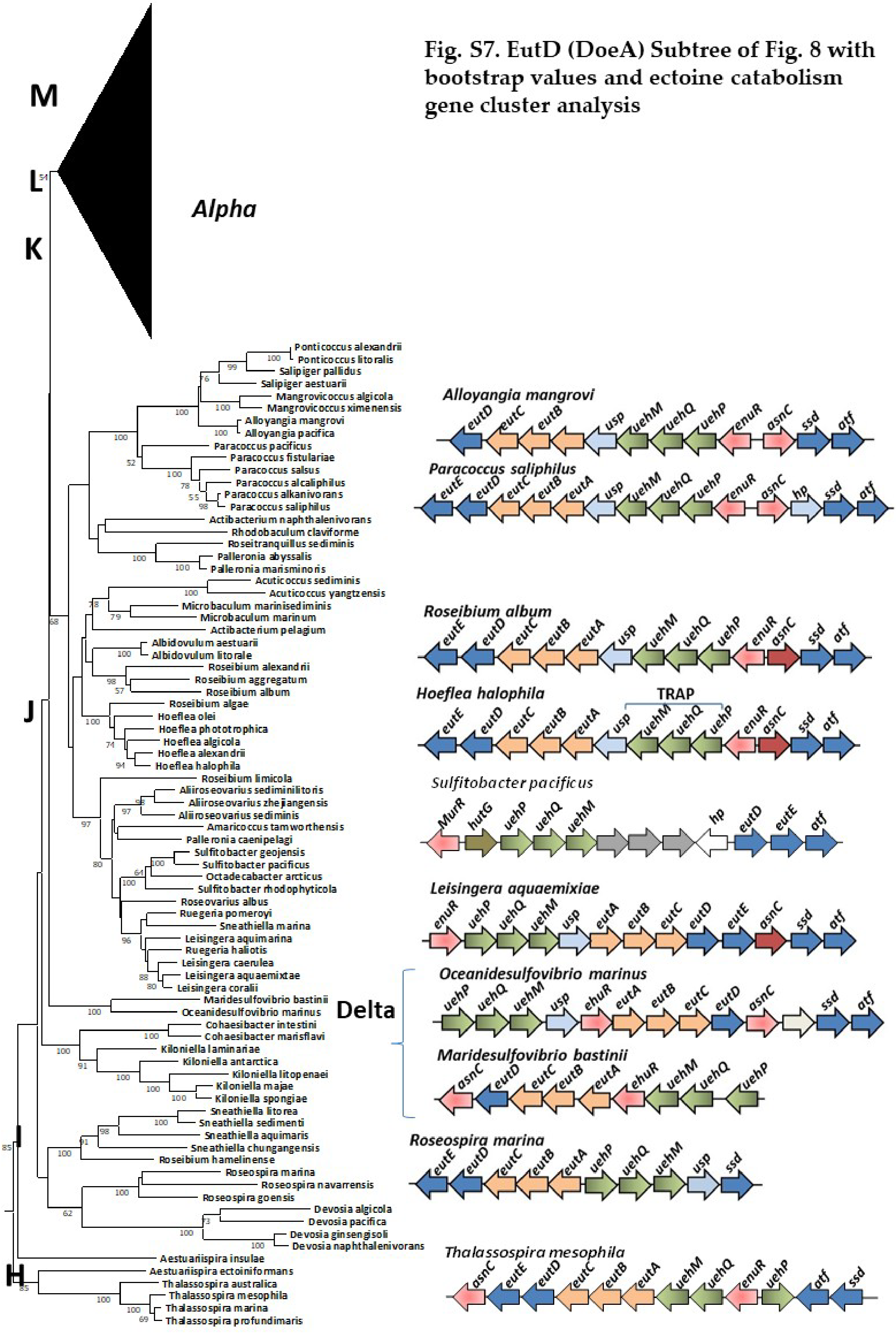


**Fig. S7. EutD (DoeA) Subtree of Fig. 8 with bootstrap values and ectoine catabolism gene cluster analysis**

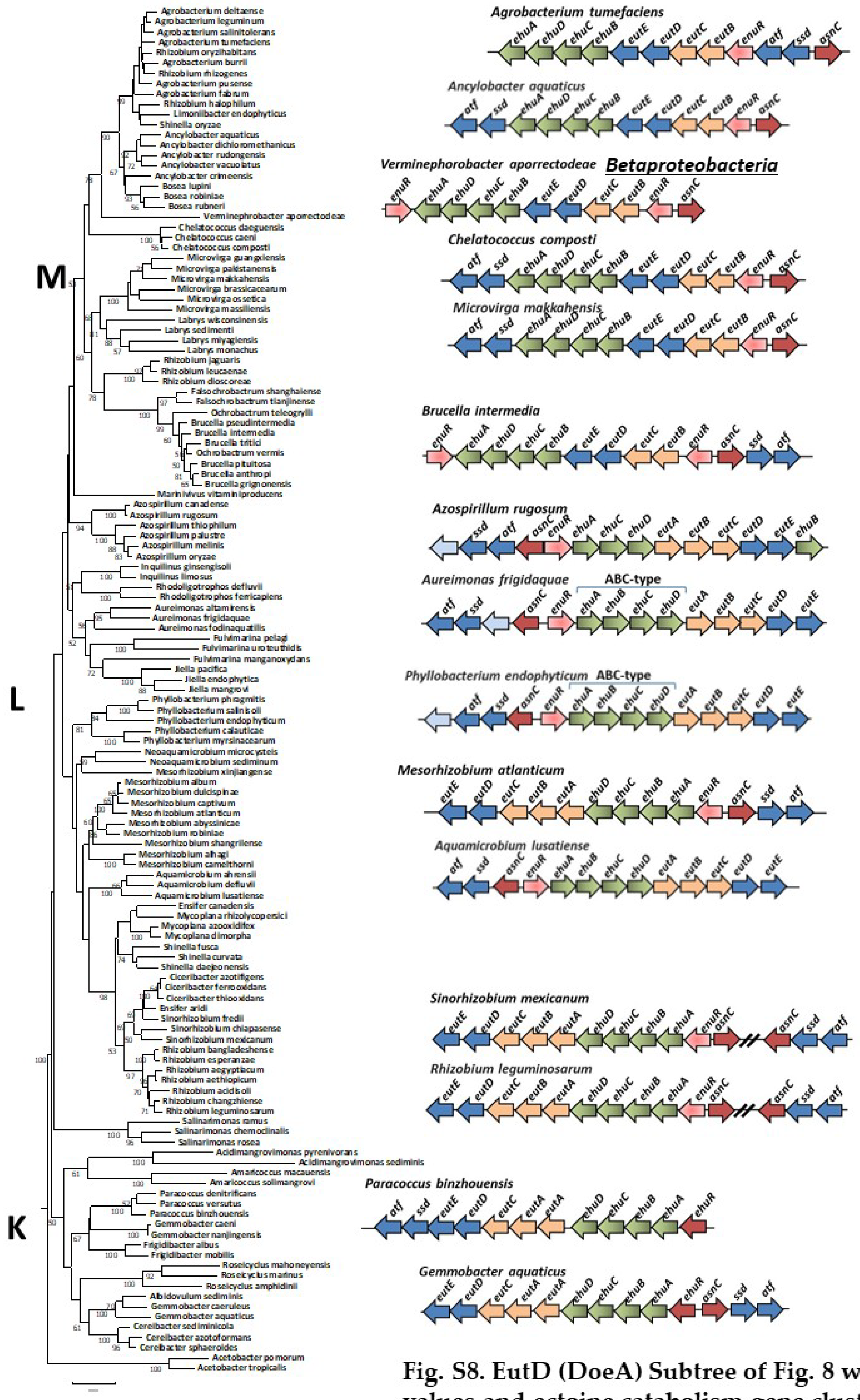


**Fig. S8. EutD (DoeA) Subtree of Fig. 8 with bootstrap values and ectoine catabolism gene cluster analysis**

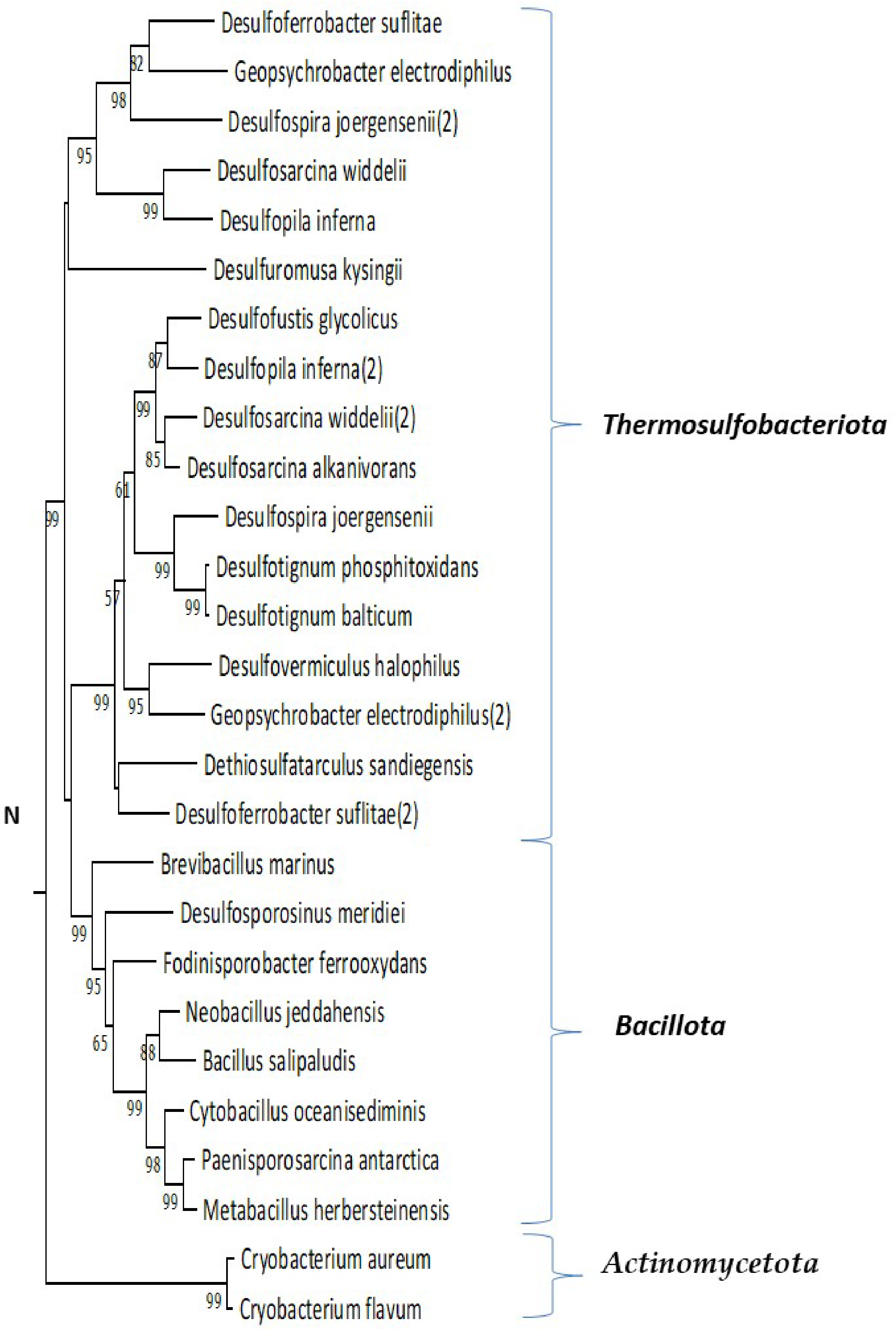


**Fig. S9. Evolutionary relationships of ectoine hydrolase EutD (DoeA) from Gram-positive bacteria.** Subtree of Fig. 8 with bootstrap values.

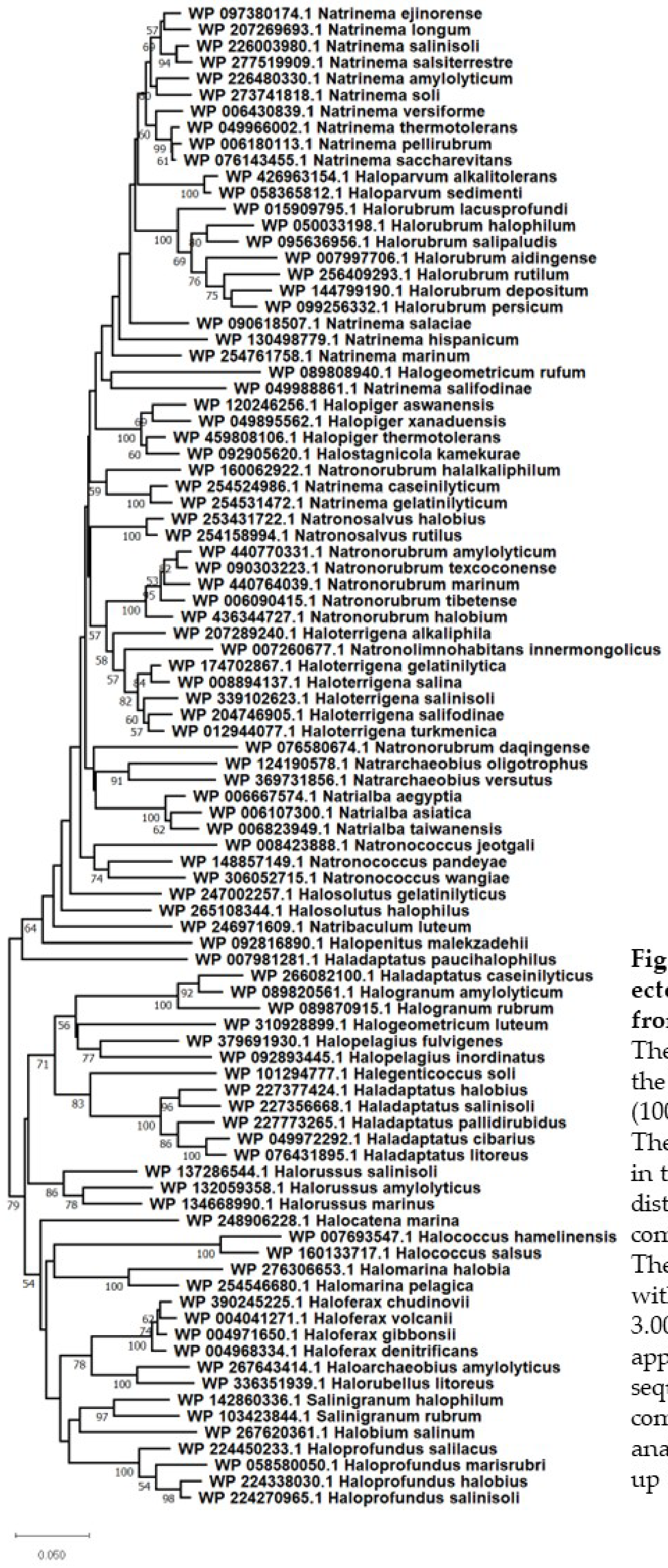


**Fig. S10. Evolutionary relationships of ectoine hydrolase EutD (DoeA) from 92 taxa from the domain *Archaea* order *Halobacteria*.** The evolutionary history was inferred using the Neighbor-Joining with the bootstrap test (100 replicates) are shown below the branches. The tree is drawn to scale, with branch lengths in the same units as those of the evolutionary distances used to infer the phylogenetic tree computed using the JTT matrix-based method. The rate variation among sites was modeled with a gamma distribution (shape parameter = 3.00). The pairwise deletion option was applied to all ambiguous positions for each sequence pair resulting in a final data set comprising 393 positions. Evolutionary analyses were conducted in MEGA12 utilizing up to 3 parallel computing threads.

**Fig. S11 A. Putative ectoine catabolism region present in Domain *Archaea*, Order *Halobacteria*. A.** *Haloprofundus halobius* genome assembly ASM2009783v1
**B.** *Halomarina pelagica* genome assembly ASM2422831v1

### A. *Haloprofundus halobius*

| | | | | | | |
|---|---|---|---|---|---|---|
| NZ_CP083667.1:173995-174972 | minus | ABC transporter permease | | LAQ74_RS17775 | 1 WP_224338024.1 | 325 |
| NZ_CP083667.1:175047-176759 | minus | ABC transporter substrate-binding protein | | LAQ74_RS17780 | 1 WP_224338026.1 | 570 |
| NZ_CP083667.1:176960-178339 | plus | **M28 family peptidase** | | LAQ74_RS17785 | 1 WP_224338028.1 | 459 |
| NZ_CP083667.1:178495-179673 | plus | **M24 family metallopeptidase** | **eutD** | LAQ74_RS17790 | 1 WP_224338030.1 | 392 |
| NZ_CP083667.1:179728-180978 | plus | glutamate dehydrogenase GdhB | **gdhB** | LAQ74_RS17795 | 1 WP_224338032.1 | 416 |
| NZ_CP083667.1:181033-184248 | minus | **FAD-binding and (Fe-S)-binding domain** | **GlcD GlpC** | LAQ74_RS17800 | 1 WP_224338034.1 | 1,071 |
| NZ_CP083667.1:184257-185228 | minus | D-2-hydroxyacid dehydrogenase | | LAQ74_RS17805 | 1 WP_224338036.1 | 323 |
| NZ_CP083667.1:185421-186710 | plus | **amidohydrolase (aminopeptidase)** | | LAQ74_RS17810 | 1 WP_224338038.1 | 429 |
| NZ_CP083667.1:186707-187984 | plus | **threonine ammonia-lyase** | **ilvA (eutB** | LAQ74_RS17815 | 1 WP_224338040.1 | 425 |
| NZ_CP083667.1:188069-189427 | minus | **aminotransferase class III-fold pyridoxal** | **AAT** | LAQ74_RS17820 | 1 WP_224338042.1 | 452 |
| NZ_CP083667.1:189580-191208 | minus | **BCCT family transporter** | **bccT** | LAQ74_RS17825 | 1 WP_224338044.1 | 542 |
| NZ_CP083667.1:191766-192413 | plus | hypothetical protein | | LAQ74_RS17830 | 1 WP_224338077.1 | 215 |
| NZ_CP083667.1:192438-192824 | minus | Rid family detoxifying hydrolase | **ridA** | LAQ74_RS17835 | 1 WP_224338046.1 | 128 |
| NZ_CP083667.1:192897-194276 | minus | **aspartate aminotransferase family protei** | **atf** | LAQ74_RS17840 | 1 WP_224338048.1 | 459 |
| NZ_CP083667.1:194384-196045 | plus | aldehyde ferredoxin oxidoreductase family protein | | LAQ74_RS17845 | 1 WP_224338050.1 | 553 |
| NZ_CP083667.1:196118-196918 | minus | **IclR family transcriptional regulator** | | LAQ74_RS17850 | 1 WP_224338052.1 | 266 |
| NZ_CP083667.1:198125-199114 | minus | PH domain-containing protein | | LAQ74_RS17855 | 1 WP_224338054.1 | 329 |
| NZ_CP083667.1:199117-200598 | minus | PH domain-containing protein | | LAQ74_RS17860 | 1 WP_224338056.1 | 493 |

### B. *Halomarina pelagica*

| | | | | | | |
|---|---|---|---|---|---|---|
| NZ_CP100455.1:488258-489361 | plus | TrmB family transcriptional regulator | | NKI68_RS20840 | 1 WP_254546666.1 | 367 |
| NZ_CP100455.1:489922-490392 | minus | helix-turn-helix domain-containing protein | | NKI68_RS20845 | 1 WP_254546667.1 | 156 |
| NZ_CP100455.1:490559-492505 | minus | archaea-specific SMC-related protein | | NKI68_RS20850 | 1 WP_254546668.1 | 648 |
| NZ_CP100455.1:492637-493251 | minus | rod-determining factor RdfA | rdfA | NKI68_RS20855 | 1 WP_254546669.1 | 204 |
| NZ_CP100455.1:493792-494973 | plus | **M24 family metallopeptidase** | pepP | NKI68_RS20860 | 1 WP_254546670.1 | 393 |
| NZ_CP100455.1:495059-495856 | plus | **IclR family transcriptional regulator** | | NKI68_RS20865 | 1 WP_254546671.1 | 265 |
| NZ_CP100455.1:495903-497564 | minus | **aldehyde ferredoxin oxidoreductase family protein** | | NKI68_RS20870 | 1 WP_254546672.1 | 553 |
| NZ_CP100455.1:497784-499163 | plus | **aspartate aminotransferase family protein** | atf | NKI68_RS20875 | 1 WP_254546673.1 | 459 |
| NZ_CP100455.1:499236-499622 | plus | **Rid family detoxifying hydrolase** | ridA | NKI68_RS20880 | 1 WP_254546674.1 | 128 |
| NZ_CP100455.1:499938-501557 | plus | **BCCT family transporter** | bccT | NKI68_RS20885 | 1 WP_254546675.1 | 539 |
| NZ_CP100455.1:501646-502998 | plus | **aminotransferase class III-fold pyridoxal phosphate-depenc** | AAT | NKI68_RS20890 | 1 WP_254546676.1 | 450 |
| NZ_CP100455.1:502991-504271 | minus | **threonine ammonia-lyase** | ilvA (eutB?) | NKI68_RS20895 | 1 WP_254546677.1 | 426 |
| NZ_CP100455.1:504268-505548 | minus | **amidohydrolase** | | NKI68_RS20900 | 1 WP_254546678.1 | 426 |
| NZ_CP100455.1:505661-506632 | plus | **D-2-hydroxyacid dehydrogenase** | | NKI68_RS20905 | 1 WP_254546679.1 | 323 |
| NZ_CP100455.1:506675-507853 | minus | **M24 family metallopeptidase** | eutD (doeA) | NKI68_RS20910 | 1 WP_254546680.1 | 392 |
| NZ_CP100455.1:508356-509585 | plus | tyrosine-type recombinase/integrase | | NKI68_RS20915 | 1 WP_254546681.1 | 409 |
| NZ_CP100455.1:510030-511400 | minus | hypothetical protein | | NKI68_RS20920 | 1 WP_254546682.1 | 456 |
| NZ_CP100455.1:511632-511796 | minus | hypothetical protein | | NKI68_RS20925 | 1 WP_254546683.1 | 54 |
| NZ_CP100455.1:512251-512766 | plus | halocyanin domain-containing protein | | NKI68_RS20930 | 1 WP_254546684.1 | 171 |
